# Spontaneous Mutations in HIV-1 Gag, protease, RT p66 in the first replication cycle and how they appear: Insights from an *in vitro* BSL2 assay on mutation rates and types

**DOI:** 10.1101/679852

**Authors:** Joshua Yi Yeo, Darius Wen-Shuo Koh, Ping Yap, Ghin-Ray Goh, Samuel Ken-En Gan

**Author notes:** Corresponding author, Antibody & Product Development Lab, BII, A*STAR, 60 Biopolis Street, #B2 Genome, Singapore 138672.

## Abstract

While drug resistant mutations in HIV-1 is largely credited to its error prone HIV-1 RT, host proteins such as deaminases may also play a role generating mutations. Many HIV-1 RT mutational *in vitro* studies utilize reporter genes (LacZ) as template, leaving out the possible contribution of HIV codon usage and gene-specific effects. To address this gap, we studied HIV-1 RT mutation rates and bias on its own Gag, protease, and RT p66 genes in an *in-vitro* selection pressure free system. We found rare clinical mutations with a general avoidance of crucial functional sites in the background mutations rates for Gag, protease and RT p66 at 4.71 x 10^−5^, 6.03 x 10^−5^, and 7.09 x 10^−5^ mutations/bp respectively. Gag and p66 genes showed a large number of ‘A to G’ hypermutations likely due to cellular adenosine deaminases. Comparisons with silently mutated p66 sequences showed an increase in mutation rates (1.88 x 10^−4^ mutations/bp) and that ‘A to G’ mutations occurred in regions reminiscent of ADAR neighbour preferences. Mutational free energies by the ‘A to G’ mutations revealed an avoidance of destabilizing effects with the natural p66 gene codon usage providing barriers to ADAR effects. Our study demonstrates the importance of studying mutation emergence in HIV genes to understand how fast drug resistance can emerge, sometimes with contributions of host deaminases, providing transferable applications to how new viral diseases and drug resistances can emerge.

## Introduction

RNA viruses have been shown to have a higher likelihood of undergoing genetic changes, leading to species jump [1] and efficient spread among humans [2]. Notably, among RNA viruses, the human immunodeficiency viruses such as HIV-1 and HIV-2 are believed to be zoonotic transmissions of the simian immunodeficiency viruses (SIV) [3,4].

Within HIV-1, the Gag and protease proteins play crucial roles in viral assembly and maturation of infectious virions [5]. Protease cleaves Gag and Pol polyproteins into functional subunits [6] and this cleaving is prevented by protease inhibitors (PI) which compete with Gag for the active site [7,8]. In emerging PI resistance, mutations on viral protease reduce affinity to PIs [8,9], and the gradual accumulation of many such resistance mutations [10-12] induced by HIV-1 RT [13,14], eventually render the PIs ineffective, and in some cases, cross-resistances to limit clinical drug selection [8]. In many cases, mutations on the substrate Gag compensate for reduced viral fitness [8,9,14-16] with paired mutations that work synergistically to protease mutations [17-21] to overcome PIs.

In the spotlight for the cause of the mutations in HIV is the error prone enzyme reverse transcriptase (RT), an asymmetric heterodimer of the p66 and p51 subunits [22]. The p66 subunit performs the key enzymatic functions of catalysing DNA polymerisation and cleaving the RNA of the RNA/DNA duplex [23,24], while the p51 subunit plays a more supportive role to p66 [25].

In the investigation of HIV mutations, in-depth analysis of RT is indispensable. However, most of the previous studies on HIV mutation rates to date utilize reporter genes such as LacZ for the template for analysis of mutations, and not HIV genes. There is thus a current gap in understanding the contribution of HIV gene-specific sequences and codon usage in mutational hotspots, as well as type of mutations and when they can emerge in the infection cycle [26]. To fill this gap, we developed an *in-vitro* based assay without translational, immune and drug selection pressures to characterize the basal HIV-1 RT mutations and biases on HIV-1 Gag, protease and RT p66 genes. As a control and further investigation to the natural codon usage, we also created a silent codon mutated variant of RT p66. Together, our findings shed light on the emergence of drug resistance mutations, the native rate, where they appear, and the associated biases and trends that would be useful for the design of future drug interventions.

## Methods

### Transfection of HIV-1 Gag, Pr and RT plasmids

HIV-1 Gag and Pr genes were PCR amplified from plasmid p8.91 [27]. HIV-1 RT p66 (GenBank: K03455.1) and codon mutated p66 genes were synthesized (BioBasic). The codon mutated p66 sequence was generated by reverse translation from the amino acid sequence for differing nucleotide (but retaining the amino acid sequence) to disrupt RNA secondary structures. The genes were then cloned separately into the pTT5 plasmid vector (YouBio), transformed into competent DH5α *E. coli* cells [28], and transfected into HEK293F cells as previously performed [29-31].

### RNA extraction and cDNA synthesis

Total RNA from cells transfected two days prior with the gene of interest were extracted using TRIzol according to manufacturer’s instructions (Invitrogen). cDNA synthesis was performed using recombinant HIV-1 RT subunits: (i) p51 (0.2475 µg) and p66 (0.2125 µg) from Sino Biological Inc (catalogue: 40244-V07E and 40244-V07E1, respectively), (ii) DNase-treated RNA, (iii) 5X RT buffer (25 mM Dithiothreitol, 375 mM KCl, 15 mM MgCl_2_, 250 mM Tris-HCl [pH 8.3]), (iv) Oligo(dT)_18_ (Thermo Scientific), (v) dNTPs, and (vi) RiboLock RNase inhibitor (ThermoFisher Scientific), in a single cycle of 25°C for 18 min, 37°C for 1 hr and 85°C for 5 min. RT negative controls were prepared without the addition of HIV-1 RT p51 and p66 subunits.

### Amplification of cDNA and TOPO cloning

PCR amplification was performed using high-fidelity Q5 Polymerase PCR (New England Biolabs) with the following primers sets: HIV-1 Gag - F (5 ‘– TAT TAG GAA TTC ATG GGT GCG AGA GCG– 3’) and R (5’ – CTG GTA AAG CTT CTA GTG GTG GTG GTG – 3’); protease - F (5’ – GCG GCC GAA TTC ATG CCT CAA ATC AC – 3’) and R (5’ – TAT AAT AAG CTT CTA GTG GTG GTG GTG –3’); RT p66 - F (5’ – ATG GCC TTG ACC TTT GCT TTA CTG – 3’) and R (5’– CTT GTC GTC ATC GTC TTT GTA GTC – 3’); and codon mutated p66 - F (5’– GCG GTG ATG GAT GGA CCA AAA GTA AA – 3’) and R (5’– CTG CGC CTA ATG ATG ATG ATG AT – 3’) as per manufacturer’s recommendations. PCR products were analysed by gel electrophoresis with GelApp [32] and purified using Gel extraction and PCR purification kits previously described [33]. Purified PCR products were cloned using Zero Blunt TOPO PCR cloning kit (Invitrogen) as per manufacturer’s protocol and transformed into in-house competent DH5α cells as previously described [28]. Transformed DH5α cells were plated and grown overnight at 37°C on LB agar plates supplemented with kanamycin (50 µg/ml). Transformants were screened using GoTaq PCR (Promega) with universal M13F (−20) forward and reverse primers prior to sequencing.

### Sequence analysis

Sequence assembly and alignment of HIV-1 sequences were performed using YAQAAT Webserver [34]. DNA2App [35] was used to analyse nucleotide and amino acid sequences. Mutations in the cDNA gene sequences were determined by multiple sequence alignments with characterised HIV-1 sequences from the Los Alamos sequence database [36]. To rule out sequencing artefacts, sequence chromatograms were analysed, with repeat re-sequencings performed for ambiguous peaks and detected mutations. The mutation rates were calculated as mutations/bp where the total number of mutations and the total nucleotide bases of the respective HIV-1 genes were compared.

A ‘Two Sample logo’ was generated to determine the underlying sequence contexts of ‘A to G’ transitions generated in our in-vitro assay for comparisons against previously reported ADAR neighbour preferences [37]. A custom script was written to locate, pool and align 4 nucleotides upstream and downstream of all adenosine mutants. Analysis of natural adenosines were also analysed to generate an ersatz set of sequences as background control (see Figure 4a for schematic representation). The mutated sites were compared against the respective ersatz background sequences using the Two Sample logo software [38] with two sample t-test without Bonferroni correction to test for significantly enriched and depleted bases within 9 nucleotides.

**Figure 1.**
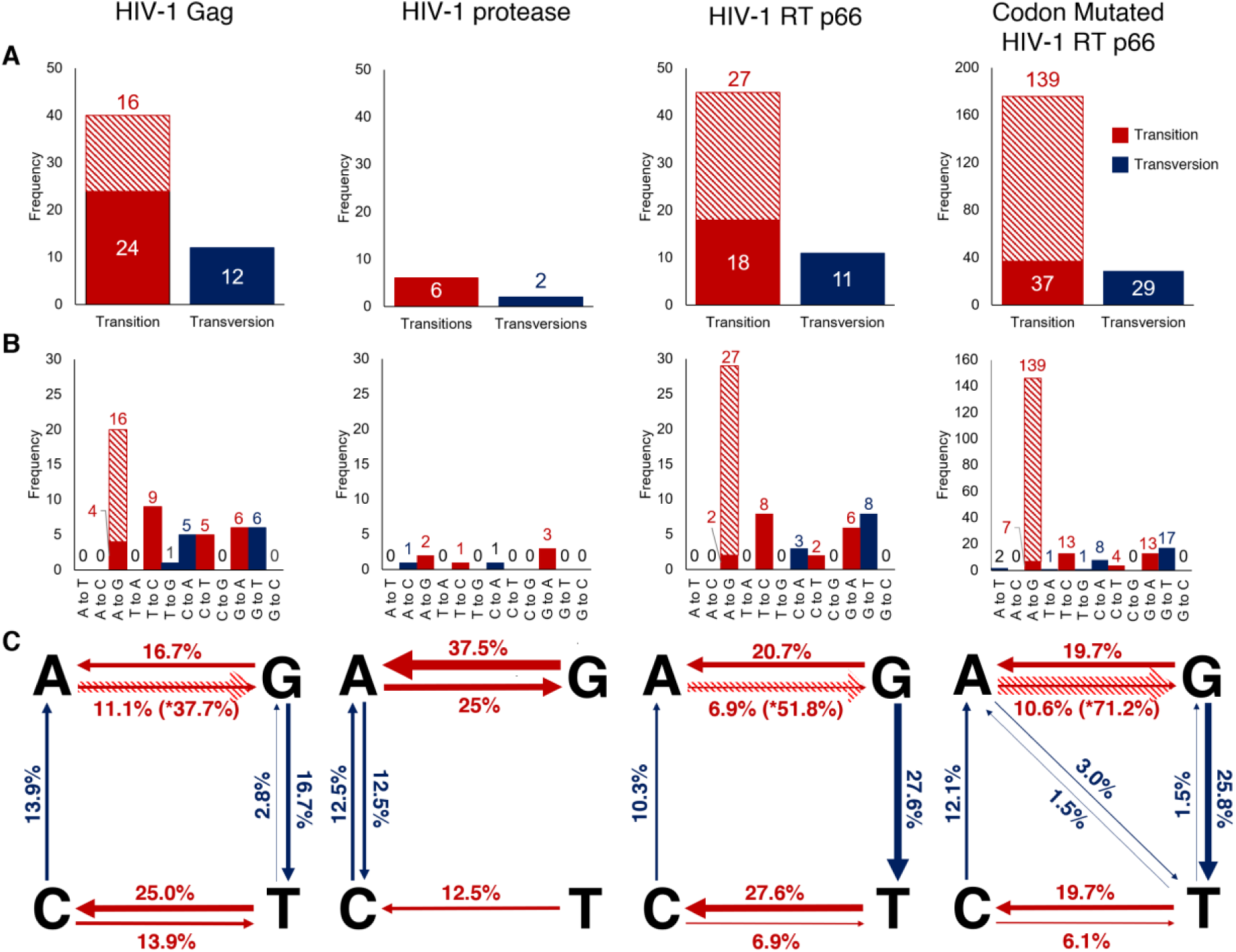
Generated HIV-1 Gag, protease, RT p66 and Codon Mutated RT p66 nucleotide substitution mutations using HIV-1 RT. (A) Bar chart of the transitions and transversions in HIV-1 Gag (n = 52), protease (n = 8), RT p66 (n = 56) and codon mutated p66 (n = 205). Transitions and transversions are shown in red and blue, respectively. ‘A to G’ hypermutations are shown separately in diagonal stripes. (B) Bar chart showing substitution mutations in HIV-1 Gag, protease, RT p66 and codon mutated p66 sequences. (C) Schematic diagram of nucleotide substitutions on the respective HIV-1 genes (expressed as a percentage).

**Figure 2.**
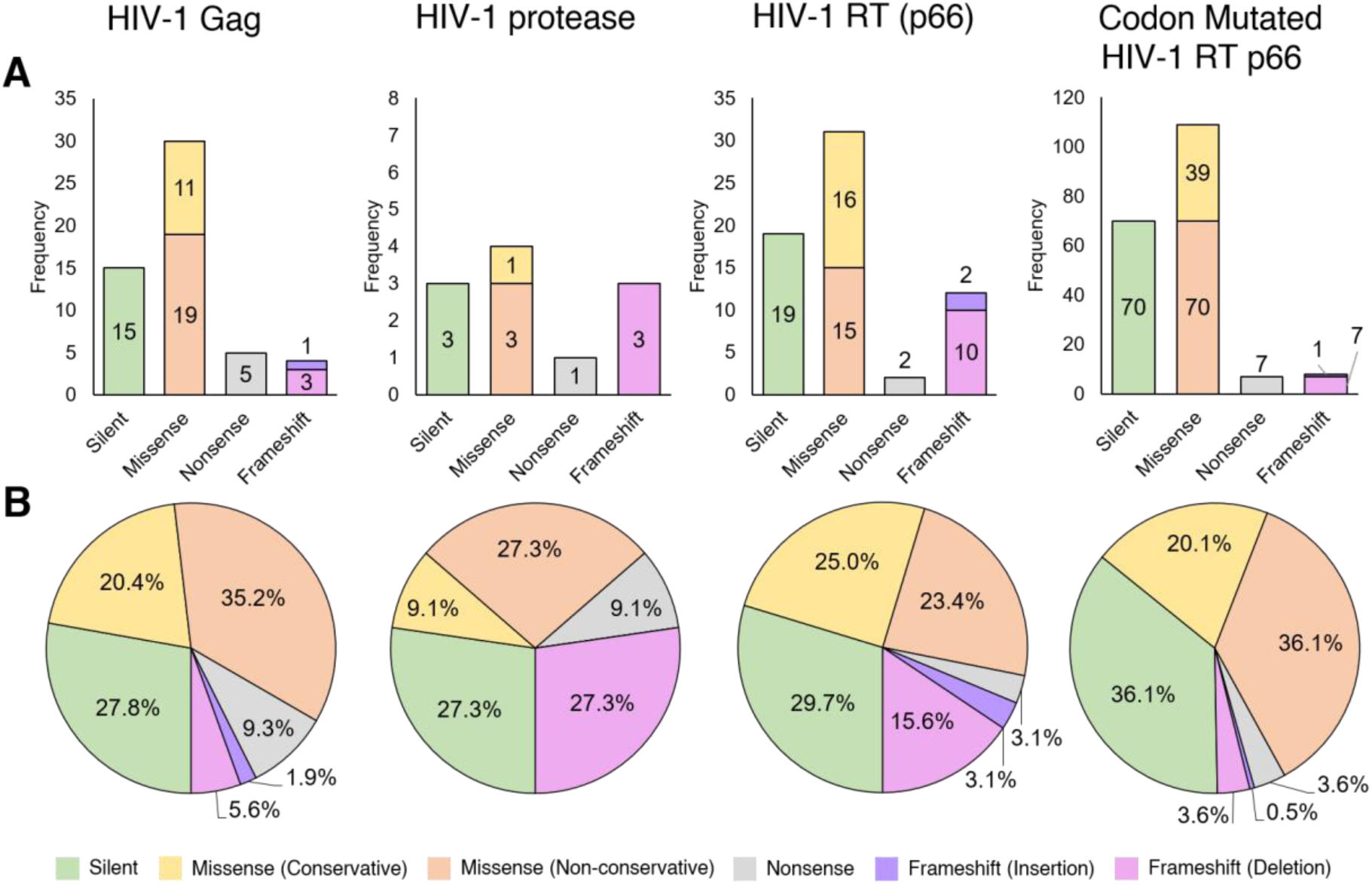
Generated amino acid mutations using HIV-1 RT. (A) Bar chart of amino acid mutations observed in HIV-1 Gag (n = 54), protease (n = 11), RT p66 (n = 64) and codon mutated RT p66 (n = 194). (B) Pie chart of the percentage proportions of the type of translation effects resulting from the mutations. For the codon mutated RT p66, two variants (20 and 80 a,b) containing deletions without frameshifts are not shown. Silent, missense (conservative), missense (non-conservative), nonsense, frameshift (insertion) and frameshift (deletion) mutations are shown in green, yellow, orange, grey, purple and pink respectively.

**Figure 3.**
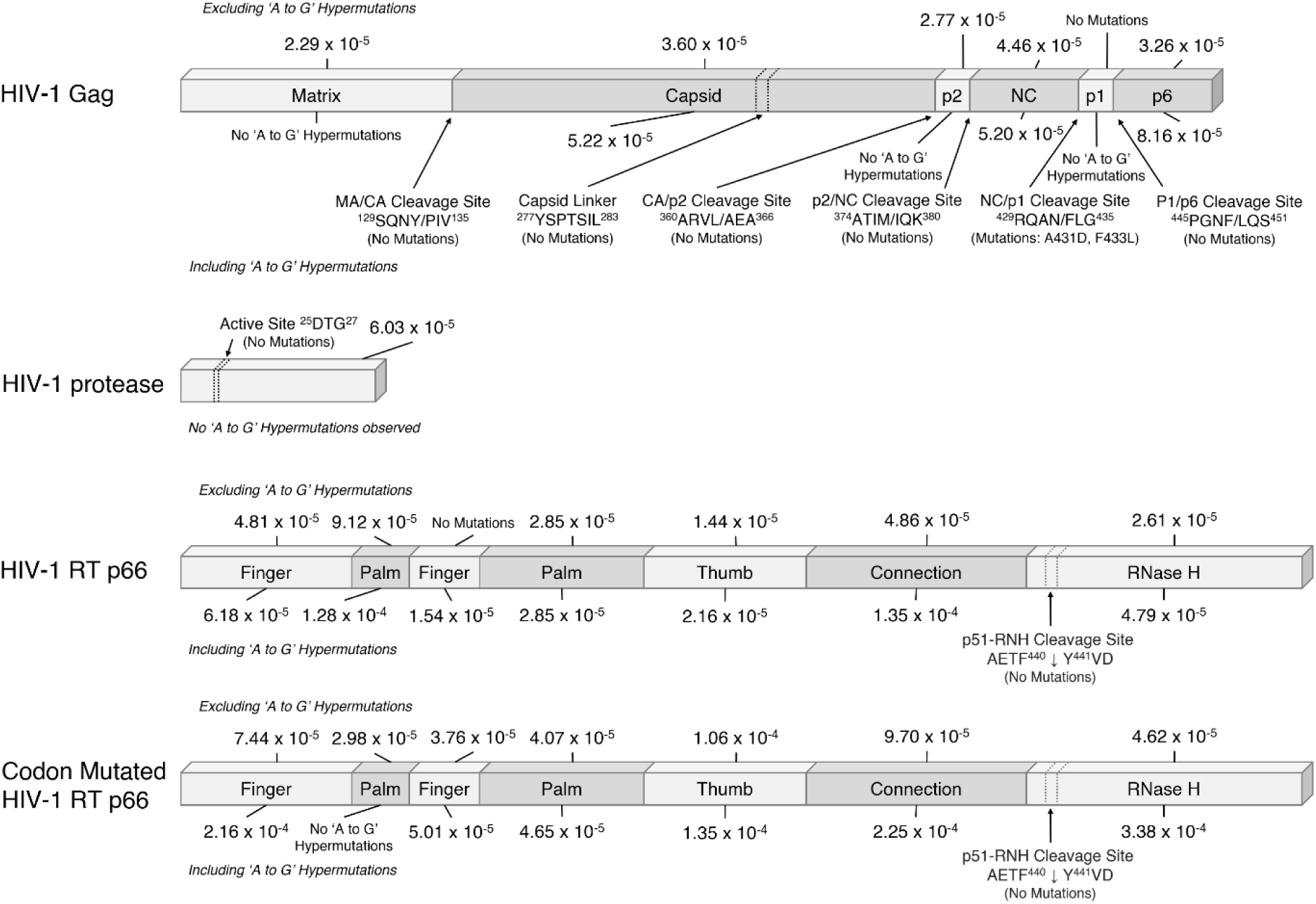
Mutation rates (mutations/bp) calculated in the domains of HIV-1 Gag, protease, RT p66 and Codon Mutated RT p66. Mutation rates calculated as the total number of mutations and the total number of nucleotide bases were counted separately excluding and including A to G hypermutations. Truncations that span across multiple domains were excluded in the calculations.

**Figure 4.**
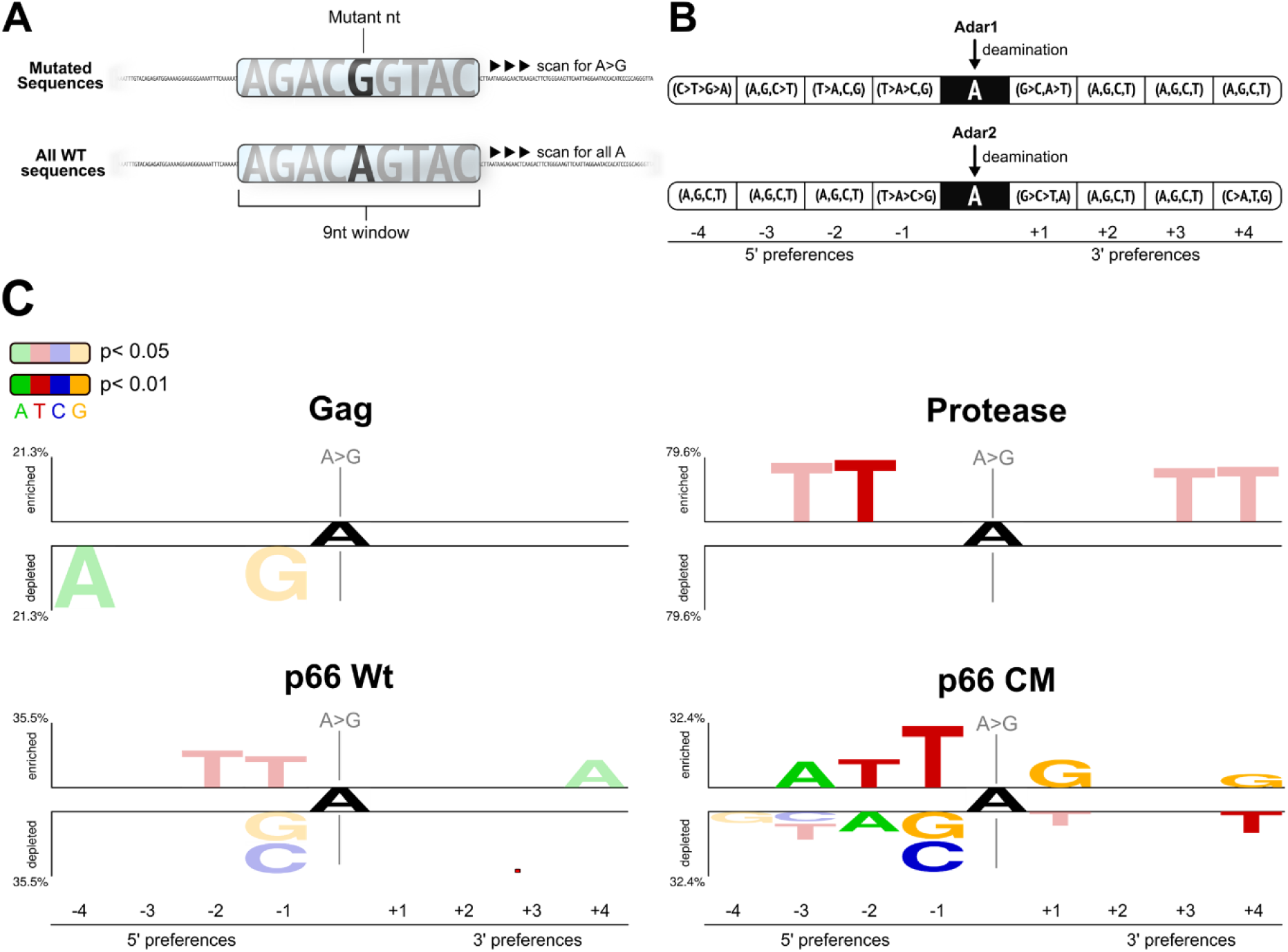
Two Sample logo analysis of HIV-1 Gag, protease, RT p66 (p66 Wt) and Codon Mutated RT (p66 CM) (A) Underlying neighbor preferences 4 nt upstream and downstream of all ‘A to G’ mutations were determined against all possible background mutations from a 9 nt sliding window. (B) A re-representation of previous reports [37] on the neighbour preferences of ADAR1 and 2 isoforms. Base preferences are shown with inequality signs, where the identity of the base did not significantly contribute to the editing of the 4 bases are delimited by commas. (C) Two Sample Logos showed enrichment and depletion of neighbouring nucleotides flanking A to G mutations. Bases were coloured opaque when p < 0.01, and translucent when p < 0.05. Statistics: two sample t-test without Bonferroni correction.

### In-silico assessment on protein thermostability using FoldX and Rosetta Cartesian_ddg

To study the mutations structurally in Gag, *in silico* mutagenesis was performed on previously modelled compact Gag structures [13] using PyMOL followed by minimization using GROMOS96 implementation in Swiss-PdbViewer. For p66, the 3T19 crystal structure was used.

Protein thermostability, free energy changes of the mutations ΔΔG_mt_ (ΔG_wild-type_ - ΔG_mutant_) were studied using FoldX5 and Rosetta Cartesian_ddg (Version : 2017.52.58848) [39]. Model structures were first relaxed and minimized using either the FoldX RepairPDB protocol or with the cartesian-space refinement relax protocol and ref2015_cart score function prior to mutagenesis. Free energy modelling was performed with the Cartesian_ddg protocol (https://www.rosettacommons.org/docs/latest/cartesian-ddG). To calculate the mutational effects on protein thermostability with FoldX, BuildModel (using default settings) was used with the numberOfRuns parameter set to 10. For Rosetta procedures, the lowest scoring structure out of 1000 cartesian-space minimization iterations were assessed to be free of serious clashes by the MolProbity server [40]. The Cartesian_ddg protocol and talaris_cart score functions were used to generate 15 models for each mutation and the average of the lowest 3 scoring mutations were averaged and multiplied by the scaling factor [39] to calculate the ΔΔG_mt_ in kcal/mols.

## Results

### Characterization of HIV-1 Gag mutant variants

In our study, we identified and classified ‘A to G’ mutations as hypermutations when multiple mutations were found on the same sequence variant given that mutations should statistically be evenly distributed.

801 HIV-1 Gag sequences were generated and calculated to show a mutation rate of 3.36 x 10^−5^ mutations/bp and at 4.71 x 10^−5^ mutations/bp when including ‘A to G’ hypermutations (see Table 1).

**Table 1:** Calculated error rates of HIV-1 RT on the respective HIV-1 genes.

| HIV-1 Gene | No. of clones | Nucleotide Length | Total no. of bases | Excluding 'A to-G' |  | Including 'A to G' |  |
| --- | --- | --- | --- | --- | --- | --- | --- |
|  |  |  |  | Hypermutations |  | Hypermutations |  |
|  |  |  |  | No. of mutations | Mutation rate (mutations/bp) | No. of mutations | Mutation rate (mutations/bp) |
| Gag | 801 | 1485 | 1,189,485 | 40 | $3.36 \times 10^{-5}$ | 56 | $4.71 \times 10^{-5}$ |
| protease | 640 | 285 | 182,400 | 11 | $6.03 \times 10^{-5}$ | - | - |
| RT p66 | 571 | 1680 | 959,280 | 41 | $4.27 \times 10^{-5}$ | 68 | $7.09 \times 10^{-5}$ |
| Codon mutated RT p66 | 700 | 1617 | 1,131,900 | 74 | $6.53 \times 10^{-5}$ | 213 | $1.88 \times 10^{-4}$ |

Mutation rates were calculated as the ratio of the total number of mutations and the total number of nucleotide bases. The rates were within the reported range of $1.8 \times 10^{-5} - 6.67 \times 10^{-4}$ as previously reviewed [26,46].

More transitions (n = 24 excluding ‘A to G’ hypermutations and n = 40 including) than transversions (n = 12) were found (Figure 1A). ‘A to G’ substitutions were found to be the most frequent (n = 20, 37.7%) when including ‘A to G’ hypermutations, but when excluded, ‘T to C’ substitutions were the most frequent (n = 9, 25.0%), followed by ‘G to A’ and ‘G to T’ (n = 6, 16.7%), ‘C to A’ and ‘C to T’ (n = 5, 13.9%), ‘A to G’ (n = 4, 11.1%), and ‘T to G’ (n = 1, 2.8%), with several substitutions types not observed (see Figure 1B and C).

Amino acid analysis showed missense mutations (n = 30, 55.6%) to occur at approximately 2 times the frequency of silent mutations (n = 15, 27.8%, see Figure 2) while nonsense mutations (n = 5, 9.3%) and frameshift mutations (n = 4, 7.5%) had lower occurrences (see Table S1 for full list of mutations).

### Characterization of HIV-1 protease mutant variants

For HIV-1 protease, 640 sequences were generated with a calculated mutation rate of 6.03 x 10^−5^ mutations/bp (see Table 1). No ‘A to G’ hypermutations were found for protease.

As with Gag, there were more transition mutations (n = 6) than transversions (n = 2) as shown in Figure 1A. ‘G to A’ substitutions were found to be the most frequent (n = 3, 37.5%), followed by ‘A to G’ (n = 2, 25%), ‘A to C’, ‘T to C’ and ‘C to A’ (n = 1, 12.5%), with several substitutions not observed (see Figures 1B and C).

Amino acid analysis showed that missense mutations (n = 4, 36.4%) occurred at approximately 1.3 times the frequency of silent mutations (n = 3, 27.3%) and frameshift mutations (n = 3, 27.3%), whereas nonsense mutations (n = 1, 9.1%) were of lower occurrences (see Figure 2, Table S2 for full list).

### Characterization of HIV-1 RT p66 mutant variants

From 571 HIV-1 RT p66 subunit sequences, we calculated a mutation rate of 4.27 x 10^−5^ mutations/bp when excluding ‘A to G’ hypermutations and 7.09 x 10^−5^ mutations/bp when including them (see Table 1).

As with Gag and protease, there were more transition mutations (n = 18 excluding ‘A to G’ hypermutations, 45 including ‘A to G’ hypermutations) than transversions (n = 11) as shown in Figure 1A. ‘A to G’ substitutions were found to be the most frequent (n = 29, 51.8%) only when including ‘A to G’ hypermutations. When excluded, ‘T to C’ and ‘G to T’ substitutions were instead the most frequent (n = 8, 27.6%), followed by ‘G to A’ (n = 6, 20.7%), ‘C to A’ (n = 4, 12.5%), and ‘A to G’ and ‘C to T’ (n = 2, 6.9%). There were several substitutions not observed (see Figure 1B and C).

Analysis after translation found similar trends for both Gag and protease where missense mutations (n = 31, 48.4%) occurred at 1.6 times the frequency of silent mutations (n = 19, 29.7%) as shown in Figure 2. Nonsense mutations (n = 2, 3.1%) and frameshift mutations (n = 12, 18.7%) were of lower occurrences (Table S3).

### Characterization of HIV-1 Codon Mutated RT p66 mutant variants

A total of 700 codon mutated p66 gave a calculated mutation rate of 6.53 x 10^−5^ mutations/bp when excluding ‘A to G’ hypermutations and 1.88 x 10^−4^ mutations/bp when including them (see Table 1).

In the same trend with Gag, protease, and wild-type p66, there were more transition mutations (n = 37 excluding ‘A to G’ hypermutations, 176 including ‘A to G’ hypermutations) than transversions (n = 29) in codon mutated p66 (Figure1A). Of the same trend with wild-type p66, ‘A to G’ substitutions were found to be the most frequent (n = 146, 71.2%) only when including ‘A to G’ hypermutations, and when excluded, ‘G to T’ substitutions were found to the most frequent (n = 17, 25.8%), followed by ‘G to A’ (n = 13, 19.7%) and ‘T to C’ (n = 13, 19.7%), ‘A to G’ (n = 7, 10.6%), ‘C to A’ (n = 8, 12.1%), ‘C to T’ (n = 4, 6.1%), ‘A to T’ (n = 2, 3.0%), and ‘T to A’ and ‘T to G’ substitutions (n = 1, 1.5%), with the absence of the following substitutions: ‘A to C’, ‘C to G’ and ‘G to C’ (see Figure. 1B and C).

Amino acid analysis showed agreement in trends with the other HIV-1 genes where missense mutations (n = 109, 56.2%) occurred approximately 1.6 times the frequency of silent mutations (n = 70, 36.1%, see Figure 2). Nonsense mutations (n = 7, 3.6%) and frameshift mutations (n = 8, 4.1%) were at lower occurrences (Table S4).

### Mutation Rates of the Domains of HIV-1 Gag, protease and RT p66

The HIV-1 genes were further analysed by their respective domains and the mutation rates calculated separately (see Figure 3). Within Gag, the overall mutation rates throughout the domains were within a narrow range of 2.29 to 4.46 x 10^−5^ mutations/bp when excluding ‘A to G’ hypermutations, but increased to 8.16 x 10^−5^ mutations/bp when including them. ‘A to G’ hypermutations were found in the capsid (CA), nucleocapsid (NC), and p6 subunits, but not in the matrix (MA), p2 and p1 domains. Mutations were not observed in the MA/CA ^129^SQNY/PIV^135^, CA/p2 ^360^ARVL/AEA^366^, p2/NC ^374^ATIM/IQK^380^, P1/p6 ^445^PGNF/LQS^451^ cleavage sites and the ^277^YSPTSIL^283^ capsid linker. However, a couple of mutations (A431D and F433L) occured in the NC/p1 ^429^RQAN/FLG^435^ cleavage site.

For HIV-1 protease, there were no ‘A to G’ hypermutations and no mutations in the active site ^25^DTG^27^. For RT p66, when excluding hypermutations, there was a range of 0 to 9.12 x 10^−5^ mutations/bp, which increased to 1.35 x 10^−4^ mutations/bp when including them. Despite the 2^nd^ finger domain being of similar length as the 1^st^ palm segment, only ‘A to G’ hypermutations were observed in the 2^nd^ finger without other mutations. No mutations were observed in the RT p51 and RNase H domain (p51-RNH) cleavage site AETF^440^↓Y^441^VD or in the catalytic triad: D110, D185 and D186. For the codon mutated p66, the mutation rate was in the range of 2.98 x 10^−5^ – 1.06 x 10^−4^ mutations/bp when excluding hypermutations and 3.38 x 10^−4^ mutations/bp when including them. Contrary to the wild-type p66, there were mutations observed in the 2^nd^ finger domain, suggesting that codon usage of the wild-type 2^nd^ finger domain may provide some barrier to mutations that are not ‘A-G’ hypermutations. Similarly, no mutations were observed in the p51-RNH cleavage site AETF^440^ ↓ Y^441^VD despite the different codon usage. Interestingly, ‘A to G’ hypermutations were not observed in the 1^st^ palm region of the codon mutated p66.

### Sequence contexts of ‘A to G’ mutations

To determine the influence of host defence deaminases, specifically ADAR in the overrepresentation of ‘A to G’ mutations in our assay as was performed on double stranded non-HIV RNA [37], we analysed the neighbouring sequences +/-4 nucleotides (nt) of all the ‘A to G’ mutations using the two-sample logo analysis (Figure 4A-C).

Analysis of the Gag showed significant depletions of ‘A’ at -4 and ‘G’ at -1 nt positions downstream of ‘A to G’ mutation sites at (p < 0.05) agreeing with known ADAR neighbour preferences [37] (see Figure 4B). Using the Two Sample Logo of multiple and single substitution mutations, there was an enrichment of ‘C’ at the +1, +4 (p < 0.05) and -4 nt (p < 0.01) positions (see Figure S1), with the former two (+1 and +4 nt positions) concurring with ADAR neighbour preferences [37] (see Figure S1 and 4B). On protease, only the ‘T’ enrichment at the -2 nt (Figure 4B) in agreement to previously reported ADAR neighbor preferences was found.

Sequence analysis of RT p66 ‘A to G’ mutations also showed signatures of ADAR [37], with enrichments of ‘T’ at the -1 over ‘G’ and ‘C’ and -2 positions (enrichment of ‘A’ at the **+**4 positions (p < 0.05).

For RT codon mutated p66 (p<0.01), there were enrichments of ‘T’ at the -1, -2 nt, enrichment of ‘A’ bases at the **+**3 nt, and enrichments of ‘G’ at the **+**1 and **+**4 nt positions (p <0.01).

For both wild-type and codon mutated p66, the large representation of multiple mutation variants masked single substitution variants with ADAR neighbour preferences found with enrichments of ‘T’ at the -1 and -2 nt positions of the 3’ end (Figure S1).

### Effects of ‘A to G’ Mutations on Protein Thermostability

The distinct observed robustness of HIV to adapt to drugs, immune pressures and the error prone nature of RTs may be attributed to the ability of viral protein structures to buffer fitness by minimizing the change in mutational free energies (ΔΔG_Mt_) [41]. When significantly destabilized (e.g. >4kcal/mol), proteins may adopt a substantially different fold or be misfolded [42]. Taking advantage of relatively large numbers of pooled ‘A to G’ mutations and other transition and transversion mutations, their distributions in ΔΔG_Mt_ were determined using protein design tools: Rosetta [39] and FoldX [43]. All the mutations were combined to calculate the change in ΔΔG_Mt_ with the threshold of significance for different distributions of ΔΔG_Mt_ at >1 kcal/mol for Rosetta [39] and ΔΔG_Mt_ >0.46 kcal/mol for FoldX [43].

The distribution of ΔΔG_Mt_ of all the mutations in Gag-p66 genes (Protease was excluded as it had no classified hypermutations) in both protein analytical tools showed that most ‘A to G’ mutations exerted little or negative changes in ΔΔG_Mt_ distributions, while other mutations typically had destabilizing effects (Figure 5A). There was also a secondary peak in the Rosetta results showing a notable distribution of ‘A to G’ transitions mutants with ΔΔG_Mt_ around a stabilizing -3.99 kcal/mol attributed to Gag (Figure 5A and 5B). From the FoldX result, there were fewer destabilizing mutations with ΔΔG_Mt_ at +5.17 kcal/mol and +7.52 kcal/mol for ‘A to G’ and other transition and transversion mutations, respectively (Figure 5A). This enforces the trend that ‘A to G’ mutations tend to elicit less ΔΔG_Mt_, especially since respective secondary peaks were also seen in Gag and the wild-type RT p66 (Figure 5B and C).

**Figure 5.**
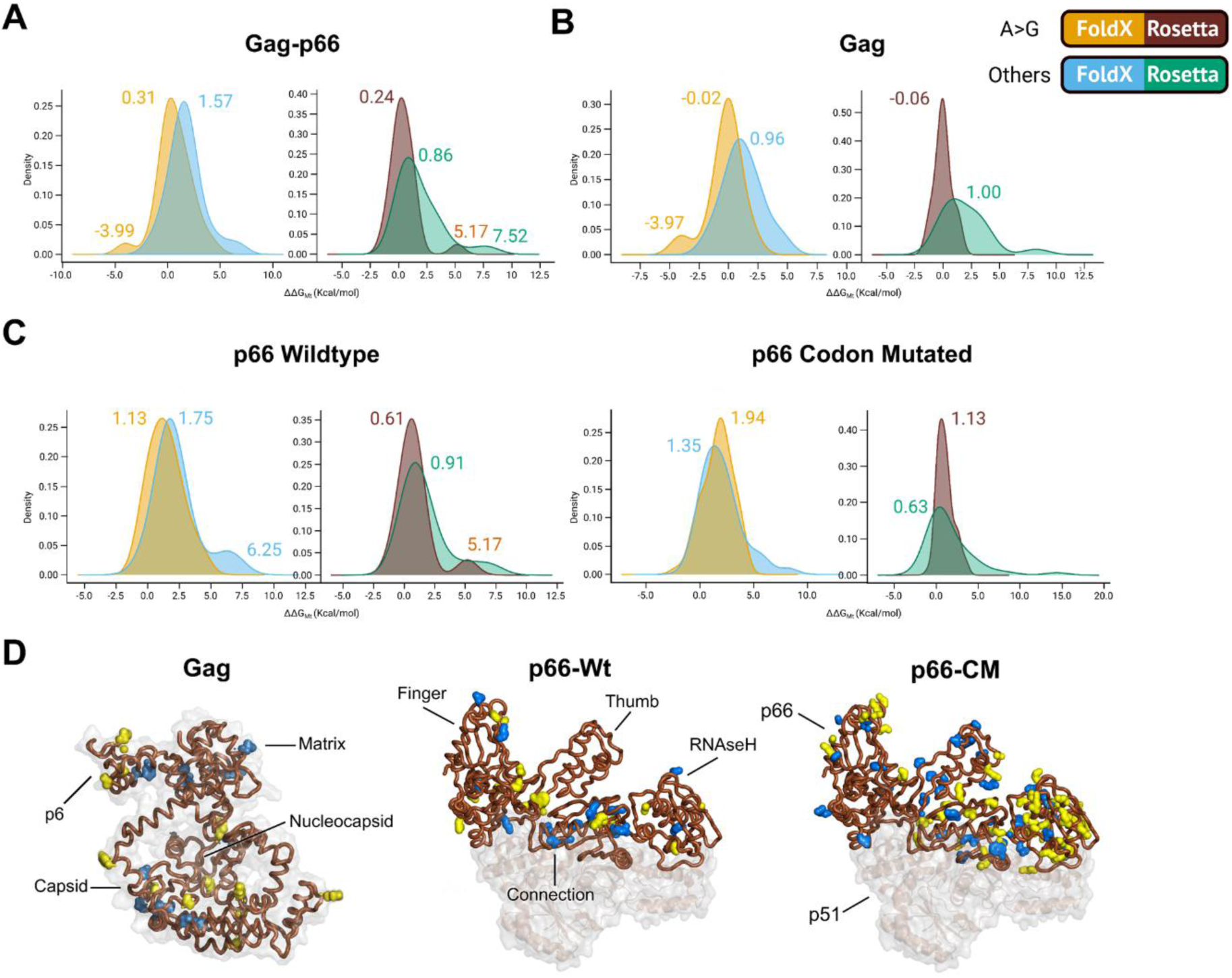
Distributions of the change in mutational free energies (ΔΔG_Mt_) of pooled mutations. Pooled ΔΔG_Mt_ of point mutations from (A) Gag-p66, (B) Gag (C) wild-type RT p66 (p66-Wt) and codon mutated RT p66 (p66-CM) genes. (D) Positions of ‘A to G’ mutations in the protein structures of Gag and p66 are shown in yellow while all other mutations are in blue. There were both ‘A to G’ and ‘other’ mutations at position 462 of p66-CM labelled blue. The C and N terminals and locations of major domains and functional protein regions are shown with labels. Peaks are numbered with corresponding ΔΔG_Mt_ values. The mutational free energies were modelled with the Rosetta Cartesian_ddg and FoldX BuildPDB protocols. ΔΔG_Mt_ values or ΔΔG_Mt_ differences between different distributions of >1 kcal for Rosetta (29) and ΔΔGMt >0.46 kcal/mol for FoldX (30) were considered to be significant. Note that Gag exists in both the Compact and Extended states and since the former precedes the later during viral assembly, the former was used.

The ‘A to G’ mutations in Gag and RT p66 genes generally had lower ΔΔG_Mt_ compared to other mutations from calculations by both protein design tools (Figure 5A-C). The FoldX results showed significant differences in the distributions of ‘A to G’ ΔΔG_Mt_ compared to other mutations in both Gag and RT p66. On the other hand, the Rosetta data did not reveal significant ΔΔG_Mt_ differences.

The results suggest that the HIV-1 genes, particularly RT p66 wild-type, were more robust to ‘A to G’ mutations, especially since ‘A to G’ mutations in codon mutated p66 were more destabilizing compared to all other mutations; with significant differences in calculations generated by FoldX.

Models of the hypermutations in Gag and wild-type p66 (Table S6) showed the average ΔΔG_Mt_ of multiple mutations as: Gag - 0.37 kcal/mol, 1.75 kcal/mol; p66 - 2.53 kcal/mol, 3.25 kcal/mol as determined using Rosetta and FoldX, respectively. These were in contrast to codon mutated p66 with an average doubling of destabilizing ΔΔG_Mt_ mutants to that of wild-type p66 at 6.48 kcal/mol, 7.00 kcal/mol for Rosetta and FoldX.

## Discussion

We set out to study the native mutation rates of HIV-1 RT on HIV-1 genes: Gag, protease and RT p66 subunit in a single replication cycle using our *in vitro* BSL2 assay that is devoid of selection pressures. In the absence of protein translational, immune, viral fitness, and drug selection pressures in our system, the observed biases are presumed to be intrinsic to HIV-RT and that of host cell factors. Although mutations can be contributed from plasmid hosts (DH5α and/or HEK293F), Q5 polymerase and even sequencing artefacts, these were highly unlikely given that the observed phenotypic mutations/bp/replication for *E. coli* and mammalian cells were estimated to be 5.4×10^−10^ and 5.0×10^−11^, respectively [44], and that Q5 polymerase is one of the most high-fidelity polymerases [45]. Admittedly, there may be a role for primer selection bias when priming the first few bases of the genes, and for this reason, these bases were excluded from analysis and calculation of mutation rates.

While numerous *in vitro* studies have reported the error rates of HIV-1 RT (as previously reviewed in [26,46]), we found only two reports that utilised HIV-1 genes as their template for analysis [47,48]. From our study, we calculated the mutation rates of HIV-1 Gag, protease, RT p66 and codon mutated p66 to be 4.71 x 10^−5^, 6.03 x 10^−5^, 7.09 x 10^−5^ and 1.88 x 10^−4^ mutations/bp (inclusive of hypermutations) respectively, within the range of 1.8 x 10^−5^ – 6.67 x 10^−4^ mutations/bp previously reported [26,46]. Across the HIV-1 genes, there was a predominance of transition mutations consistent across most phyla. This transition bias was hypothesized to be for the conservation of protein functions [49-51]. In our assay, we observed missense mutations to occur at the highest frequency, followed by silent mutations, frameshifts and nonsense. These observations are in agreement with the in-built genetic code probabilities [52] to maintain protein translatability, and avoidance of detrimental effects from frameshifts and nonsense mutations.

The use of different HIV-1 gene templates also elicited varying mutation rates, type of mutations and mutational biases. We noted a general absence of ‘A – T’ or ‘C – G’ mutations in the wild-type HIV genes studied, detecting these mutations only in the codon mutated p66 (see Figure 1), suggesting influences that are inbuilt in the sequences and codon usage. This is clearly seen when the codon mutated p66 mutation rate was 2.7 folds higher than the wild-type HIV-1 RT p66. Interestingly, the RNase H domain of the codon mutated HIV-1 RT p66 was also found to have a 7.1 fold increase (3.38 x 10^−4^ mutations/bp) when compared to the wild-type (4.79 x 10^−5^ mutations/bp), possibly due to RNA secondary structural protection/susceptibility [37,53,54].

The increase in ‘A to G’ hypermutations were observed in HIV-1 Gag, RT p66 and codon mutated RT p66, but not in protease. Possible explanations may be due to the short nucleotide length of protease, thereby avoiding the ‘A to G’ mutational effects caused by host adenosine deaminases such as double-stranded RNA-specific adenosine deaminase (ADAR). ADAR editing (by either ADAR1 and ADAR2) have been suggested to influence such cell based *in vitro* RT-fidelity assays [55]. ADAR deaminates ‘A’ residues to inosine (I) residues (read as ‘G’ by ribosomes due to structural homology), leading to ‘C’ mis-incorporations in the plus strand DNA after reverse transcription, resulting in ‘G’ residues being incorporated during second (minus) stand synthesis [56,57]. ADAR deaminases are believed to be host defence mechanism proteins against viral infections by exerting significant changes on the genomes of such RNA based lifeforms [58-60], exhibiting both pro-viral and anti-viral activities [61-64].

We also found lower occurrences of ADAR neighbour preferences bases in protease (Figure 4) which may explain the lower occurrences of PI resistance in clinical studies [65], alongside better potency than reverse transcriptase inhibitors (RTIs) [66].

The susceptibility of Gag, p66 and codon mutated p66 to ‘A to G’ hypermutations would result in an accumulation of G bases, that would lead to a translational bias towards glycine, arginine, valine and alanine [52]. This will in turn reduce the occurrences of phenylalanine, isoleucine, tyrosine, histidine and asparagine in these genes. For HIV-1 protease, the mutational bias towards ‘A’ and ‘C’ accumulation, which is also the case when ‘A to G’ hypermutations are not considered for Gag and both p66 genes, would lead to translational bias in the ‘A’ accumulation towards threonine and lysine, isoleucine, asparagine, arginine and stop codons, while reducing phenylalanine, tryptophan and cysteine occurrences. Similarly, the accumulation of ‘C’ would lead to proline, serine, leucine, threonine, alanine and arginine while reducing methionine, lysine, glutamic acid, tryptophan, and more importantly, stop codons. It is expected that higher mutations rates can push the virus towards lethal mutagenesis, allowing a possible therapeutic intervention to exploit the ADAR neighbour preference bases flanking target sites.

Structurally, the ‘A to G’ transitions elicit less destabilization effects in ΔΔG_Mt_ within Gag and wild-type p66. Gag was found to be more robust compared to p66 with its ΔΔG_Mt_ being generally neutral (Rosetta data showed distributions of significantly stabilizing mutations that could have acted as epistatic buffers against destabilizing ones), although this may require confirmation using cleaved Gag models [67]. On the other hand, codon mutated p66 was less resilient to ‘A to G’ changes compared to wild-type p66, hinting that RNA viral protein do not just minimize ΔΔG_Mt_ at the structural level as suggested by Tawfik, Berezovsky and colleagues [41], but also at the nucleotide sequence level. With multiple ‘A to G’ substitutions by ADAR hyper-editing, the average ΔΔG_Mt_ of all p66 mutations (and also Gag) were half that of the codon mutated p66. Lower rates of ‘A to G’ transition occurred in the wild-type p66 compared to its codon mutated counterpart. Such observations clearly demonstrate that the codon usage of the natural wild-type p66 (and possibly Gag) gene offered some protection against ADAR.

Despite the lack of selection pressure towards functional proteins, we did not find mutations in active sites of the enzymes (protease and p66) nor in the cleavage sites of Gag, with the exception of A431D and F433L in the NC/p1 cleavage site ^429^RQAN/FLG^435^ of Gag. The detection of non-cleavage mutations in Gag show that such compensation mutations would emerge early in the infection process even in the absence of drugs. While mimiking a single replication cycle, we found known clinical drug resistance mutations (see Table S5 for Gag, protease, and p66). In Gag, the previously reported rare transient mutation E17K (in the matrix), implicated in cytotoxic T lymphocyte (CTL) immune evasion resistance [68,69] was detected. In protease, K70T, a minority mutation associated with resistance to PIs [70] was also found. In p66, a known polymorphic mutation K103R (in the palm domain), which when combined with V179D (not found in our assay), reduced susceptibility to non-nucleoside reverse transcriptase inhibitors (NNRTIs): nevirapine (NVP) and efavirenz (EFV) by about 15 folds [71]. Interestingly. the observed K103R originated from an ‘A to G’ hypermutation event, specifically from “AAA” to “AGA” suggesting that drug resistance mutations can be induced by ADAR.

Though in much reduced frequency, some mutations were also found in crucial functional sites. In Gag, P222L occurred in the cyclophilin A (CyPA) binding site (along with G221), potentially affecting the binding of the capsid to CyPA as previously reported for P222A [72]. Given that the probability of P (CCT) to L (CTT) is 0.33 in having a T substitution in the 2^nd^ codon position, and that P (CCT) to A (GCT) is also 0.33 in having the first C mutated to a G, there were equal probabilities. However, when considering that our Gag mutation analysis did not show a transition mutation link between C to G (Figure 1), the clinical mutation is a rare-occurrence (under additional selection pressures like drug) or a result of two or more mutation steps. Mutations A431D in the nucleocapsid and P453T in the p6 domain were also observed, with reported clinical drug resistance counterparts as A431V and a L449F/P453T pair [73]. In p66, the substitution F61S was observed, implicated in strand synthesis with F61Y/L/W altering activity [74]. Similarly, we observed P95L, in which P95 was reported to be at the dimerization interface for the formation of the bottom of NNRTI pocket [75], and a proposed target amino acid in NNRTI design together with N137 and P140 [76]. Although P95 was reported to be in a highly conserved location [75], it was present in our assay with a much limited sample size compared to the virions generated daily in an infected patient.

The occurrence of previously reported *in vitro* and clinical mutations in our assay show that these mutations occurred very early in infection without immune surveillance and drug selection pressures, possibly contributed by HIV gene specific biases (Figure 1). The general avoidance of inducing mutations at crucial sites at the nucleotide stage and protective effects in natural nucleotide codon usage (compared to p66 codon mutated) to minimize changes in protein ΔΔG_Mt_, seems to suggest in-built sequence barriers of self-preservation. Of clinical relevance is that these mutations occurred within our mimic of a single cycle of replication, which when taking into account that HIV-1 generates approximately 10^9^ virions per day in an infected individual [77] drug resistance virions would have been made even within the first day of infection.

In the absence of selection pressures, mutations that are selected against can be better detected, allowing for a more comprehensive analysis of the HIV-1 RT bias that can be missed in multiplexing methods. Given that many HIV proteins can function in intense drug/immune selection environments with significant reduced activity [78,79], an all-out campaign against HIV could involve inhibitors against low-frequency occurring drug resistance mutations. This pushes the HIV infection towards Muller’s ratchet [80-82] or by increasing viral mutation rate beyond pass a critical threshold towards lethal mutagenesis for error catastrophe [83,84]. Since HIV-1 replicates at the edge of this critical threshold [85], a slight increase in the mutation rate may be sufficient to bring about error catastrophe, which have been successfully induced by mutagenic nucleoside analogues [86-88].

Coupling with *in silico* tools, such as those incorporating probability mutational change, it may be possible to identify mutational hotspots. As it was previously shown that protease and RT drug cross resistance have a structural basis governed by drug resistance mutations [89,90], the bias of restricting specific amino acid changes by the absence of A-T, C-G mutation occurrences can be exploited. However, such an approach will require an in-depth understanding of HIV-RT mutations that are selected against at protein functional levels.

Through the mutational biases and mutation rate of HIV-1 RT, it is possible to calculate the mutational events leading to the zoonotic transmission of SIV to HIV, opening up surveillance of emerging viral threats [2], especially given that RNA viruses are the most likely to species jump [1].

In conclusion, we have established an assay and characterised HIV-1 RT mutations on HIV-1 Gag, protease and RT p66 in a safe BSL2 environment, allowing for insights to the mutational bias and mutation rate of HIV in the absence of biological selection pressures. Through our analysis, we found effects of ADAR in increasing the genetic diversity beyond RT to generate rare and documented clinical drug resistance mutations within a single replication cycle and thereby very early in infection. The assay can provide deeper insights relevant for drug and vaccine development and be applied for horizontal understanding to other viruses.

## Acknowledgments

We thank Chinh Tran-To Su, Wai-Heng Lua and Wei-Li Ling for useful comments and discussion.

## Author Contributions

JYY, PY, and GRG performed the *in vitro* experiments. JYY, PY, GRG, and DWSK analysed the results. DWSK performed the *in silico* analysis. JYY, PY, GRG, DWSK, and SKEG discussed and drafted the manuscript. SKEG conceived the idea and supervised the whole project. All authors read and approved the manuscript.

