## Supplementary Data for "Spontaneous Mutations in HIV-1 Gag, protease, RT p66 in the first replication cycle and how they appear: Insights from an *in vitro* BSL2 assay on mutation rates and types"

### SUPPLEMENTARY INFORMATION

**Table S1 HIV-1 Gag nucleotide and amino acid mutations.** Free energy values $\Delta\Delta G_{Mt}$ (kcal / mol) were rounded to 2 decimal places.

| Variant | Nucleotide Position | Initial Codon | Mutated Codon | Nucleotide Mutation | Amino Acid Position | Initial Amino Acid | Mutated Amino Acid | Amino Acid Mutation | Rosetta Cartesian_ddg $\Delta\Delta G_{Mt}$ (kcal / mol) | FoldX BuildPDB $\Delta\Delta G_{Mt}$ (kcal / mol) | Type of Mutation | | Domain |
|---|---|---|---|---|---|---|---|---|---|---|---|---|---|
| 1 | 49 | GAA | AAA | G49A | 17 | E (Glutamic Acid) | K (Lysine) | E17K | -0.53 | -0.19 | Missense | Non-conservative | MA |
| 1 | 759 | AGT | AGC | T759C | 253 | S (Serine) | S (Serine) | - | - | - | Silent | | CA |
| 2 | 574 | GGG | AGG | G574A | 192 | G (Glycine) | R (Arginine) | G192R | -0.79 | 0.56 | Missense | Non-conservative | CA |
| 3 | 722 | AGT | ATT | G722T | 241 | S (Serine) | I (Isoleucine) | S241I | -1.81 | 3.00 | Missense | Non-conservative | CA |
| 4A,B | 1297 | TTT | CTT | T1297C | 433 | F (Phenylalanine) | L (Leucine) | F433L | -0.27 | 2.98 | Missense | Conservative | NC |
| 5 | 812 | AAT | AGT | A812G | 271 | N (Asparagine) | S (Serine) | N271S | 0.99 | -0.20 | Missense | Conservative | CA |
| 6 | 1451 | TTT | TCT | T1451C | 484 | F (Phenylalanine) | S (Serine) | F484S | 4.65 | 0.09 | Missense | Non-conservative | P6 |
| 7 | 371 | CAC | CGC | A371G | 124 | H (Histidine) | R (Arginine) | H124R | -0.17 | 0.00 | Missense | Conservative | MA |
| 7 | 869 | AAG | AGG | A869G | 290 | K (Lysine) | R (Arginine) | K290R | -1.53 | -0.20 | Missense | Conservative | CA |
| 8 | 1253 | AAG | AGG | A1253G | 418 | K (Lysine) | R (Arginine) | K418R | 0.07 | -0.36 | Missense | Conservative | NC |
| 8 | 715 | ACT | GCT | A715G | 239 | T (Threonine) | A (Alanine) | T239A | 0.26 | 3.75 | Missense | Non-conservative | CA |
| 9A,B | 1359 | CCA | CCG | A1359G | 453 | P (Proline) | P (Proline) | - | - | - | Silent | | P6 |
| 9A,B | 605, 606 | AAA | AGG | A605G, A606G | 202 | K (Lysine) | R (Arginine) | K202R | 0.06 | 0.01 | Missense | Conservative | CA |
| 9A,B | 610 | ACC | GCC | A610G | 204 | T (Threonine) | A (Alanine) | T204A | 0.29 | 1.27 | Missense | Non-conservative | CA |
| 9A,B | 667 | ATT | GTT | A667G | 223 | I (Isoleucine) | V (Valine) | I223V | -0.11 | 0.11 | Missense | Conservative | CA |
| 9A,B | 1386 | AGC | AGA | C1386A | 462 | S (Serine) | R (Arginine) | S462R | 1.92 | 2.86 | Missense | Non-conservative | P6 |
| 9A,B | 640 | AGA | GGA | A640G | 214 | R (Arginine) | G (Glycine) | R214G | 1.93 | -0.36 | Missense | Non-conservative | CA |
| 10 | 779 | - | - | - | - | - | - | - | - | - | Frameshift | Insertion | CA |
| 11A,B | 43 | CGA | TGA | C43T | 15 | R (Arginine) | Stop | - | - | -0.20 | Nonsense | | MA |
| 12 | 603 | TTA | TTG | A603G | 201 | L (Leucine) | L (Leucine) | - | - | - | Silent | | CA |

| | | | | | | | | | | | | | |
|---|---|---|---|---|---|---|---|---|---|---|---|---|---|
| 12 | 606 | AAA | AAG | A606G | 202 | K (Lysine) | K (Lysine) | - | - | - | Silent | | CA |
| 12 | 630 | GCA | GCG | A630G | 210 | A (Alanine) | A (Alanine) | - | - | 0.30 | Silent | | CA |
| 13 | 964 | TTG | CTG | T964C | 322 | L (Leucine) | L (Leucine) | - | - | - | Silent | | CA |
| 14 | 1116 | AAT | AAC | T1116C | 372 | N (Asparagine) | N (Asparagine) | - | - | 1.27 | Silent | | SP2 |
| 15 | 1281 | ACT | ACC | T1281C | 427 | T (Threonine) | T (Threonine) | - | - | - | Silent | | NC |
| 16 | 665 | CCT | CTT | C665T | 222 | P (Proline) | L (Leucine) | P222L | 0.76 | 0.20 | Missense | Conservative | CA |
| 16 | 124 | GAA | AAA | G124A | 42 | E (Glutamic Acid) | K (Lysine) | E42K | 2.4 | 0.74 | Missense | Non-conservative | MA |
| 17A,B | 184 | GGA | TGA | G184T | 62 | G (Glycine) | Stop | - | - | 2.86 | Nonsense | | MA |
| 18 | 316 | GAA | TAA | G316T | 106 | E (Glutamic Acid) | Stop | - | - | - | Nonsense | | MA |
| 19A,B | 443 | TCA | TAA | C443A | 148 | S (Serine) | Stop | - | - | 1.23 | Nonsense | | CA |
| 20 | 454 | TTA | GTA | T454G | 152 | L (Leucine) | V (Valine) | L152V | 2.7 | 1.38 | Missense | Conservative | CA |
| 21 | 525 | TTA | TTG | A525G | 175 | L (Leucine) | L (Leucine) | - | - | - | Silent | | CA |
| 22A,B | 733 | GAA | AAA | G733A | 245 | E (Glutamic Acid) | K (Lysine) | E245K | 2.15 | - | Missense | Non-conservative | CA |
| 23A,B | 834 | AGC | AGT | C834T | 278 | S (Serine) | S (Serine) | - | - | - | Silent | | CA |
| 24 | 921 | GAG | GAA | G921A | 307 | E (Glutamic Acid) | E (Glutamic Acid) | - | - | - | Silent | | CA |
| 25 | 1021 | GCG | TCG | G1021T | 341 | A (Alanine) | S (Serine) | A341S | 0.39 | - | Missense | Non-conservative | CA |
| 25 | 1186 | GGC | AGC | G1186A | 396 | G (Glycine) | S (Serine) | G396S | 0.95 | - | Missense | Non-conservative | NC |
| 26 | 1050 | TGT | TGC | T1050C | 350 | C (Cystine) | C (Cystine) | - | - | - | Silent | | CA |
| 27 | 1071 | GGC | GGT | C1071T | 357 | G (Glycine) | G (Glycine) | - | - | - | Silent | | CA |
| 27 | 445 | CCT | ACT | C445A | 149 | P (Proline) | T (Threonine) | P149T | 0.78 | -0.27 | Missense | Non-conservative | CA |
| 28 | 1146 | AAT | AAC | T1146C | 382 | N (Asparagine) | N (Asparagine) | - | - | - | Silent | | NC |
| 29 | 1250 | GGA | GTA | G1250T | 417 | G (Glycine) | V (Valine) | G417V | 3.82 | 0.74 | Missense | Conservative | NC |
| 30A,B | 1292 | GCT | GAT | C1292A | 431 | A (Alanine) | D (Aspartic Acid) | A431D | 0.87 | 0.73 | Missense | Non-conservative | NC |
| 31 | 1357 | CCA | ACA | C1357A | 453 | P (Proline) | T (Threonine) | P453T | 2.15 | - | Missense | Non-conservative | P6 |
| 32A,B,C,D,E | 1360 | GAG | TAG | G1360T | 454 | E (Glutamic Acid) | Stop | - | - | - | Nonsense | | P6 |
| 33A,B,C | 1408 | ACA | GCA | A1408G | 470 | T (Threonine) | A (Alanine) | T470A | -0.84 | - | Missense | Non-conservative | P6 |
| 33A,B,C | 1411 | ACT | GCT | A1411G | 471 | T (Threonine) | A (Alanine) | T471A | -4.03 | 1.38 | Missense | Non-conservative | P6 |
| 34 | 673 | CCA | TCA | C673T | 225 | P (Proline) | S (Serine) | P225S | 0.99 | 1.41 | Missense | Non-conservative | CA |
| 35 | 137 | GTT | GCT | T137C | 46 | V (Valine) | A (Alanine) | V46A | 0.77 | - | Missense | Conservative | MA |

| | | | | | | | | | | | | | |
|---|---|---|---|---|---|---|---|---|---|---|---|---|---|
| 36A,B,C,D, E | 673 – 1369 | - | - | - | - | - | - | - | - | 8.22 | Frameshift | Deletion | CA TO P6 |
| 37 | 673 – 1370 | - | - | - | - | - | - | - | - | - | Frameshift | Deletion | CA TO P6 |
| 38 | 1497 | TCA | TCG | A1497G | 499 | S (Serine) | S (Serine) | - | | - | Silent | | P6 |
| 38 | 1499, 1500 | CAA | CGG | A1499G, A1500G | 500 | Q (Glutamine) | R (Arginine) | Q500R | -0.32 | - | Missense | Non-conservative | P6 |
| 39 | 578 | - | - | - | - | - | - | - | - | 0.39 | Frameshift | Deletion | CA |

### Table S2 HIV-1 Protease nucleotide and amino acid mutations.

| Variant | Nucleotide Position | Initial Codon | Mutated Codon | Nucleotide Mutation | Amino Acid Position | Initial Amino Acid | Mutated Amino Acid | Amino Acid Mutation | Type of Mutation | |
|---|---|---|---|---|---|---|---|---|---|---|
| 1 | 291 | TTA | TTG | A291G | 97 | L (Leucine) | L (Leucine) | - | Silent | |
| 2 | 44 | GTA | GCA | T44C | 15 | V (Valine) | A (Alanine) | V15A | Missense | Conservative |
| 3 | 292 | AAT | GAT | A292G | 98 | N (Asparagine) | D (Aspartic Acid) | N98D | Missense | Non-conservative |
| 4 | 270 | TTG | TTA | G270A | 90 | L (Leucine) | L (Leucine) | - | Silent | |
| 5 | 236 | CCT | CAT | C236A | 79 | P (Proline) | H (Histidine) | P79H | Missense | Non-conservative |
| 6 | 209 | AAG | ACG | A209C | 70 | K (Lysine) | T (Threonine) | K70T | Missense | Non-conservative |
| 7 | 126 | TGG | TGA | G126A | 42 | W (Tryptophan) | Stop | - | Nonsense | |
| 8 | 114 | TTG | TTA | G114A | 38 | L (Leucine) | L (Leucine) | - | Silent | |
| 9 | 136 | - | - | - | - | - | - | - | Frameshift | Deletion |
| 10 | 257 – 286 | - | - | - | - | - | - | - | Frameshift | Deletion |
| 11A-E | 297 | - | - | - | - | - | - | - | Frameshift | Deletion |

**Table S3 HIV-1 RT p66 nucleotide and amino acid mutations.** Free energy values $\Delta\Delta G_{Mt}$ (kcal / mol) were rounded to 2 decimal places.

| Variant | Nucleotide position | Initial codon | Mutated codon | Nucleotide Mutation | Amino Acid Position | Initial Amino Acid | Mutated Amino Acid | Amino Acid Mutation | Rosetta Cartesian_ddg $\Delta\Delta G_{Mt}$ (kcal / mol) | FoldX BuildPDB $\Delta\Delta G_{Mt}$ (kcal / mol) | Type of mutation | | Domain |
|---|---|---|---|---|---|---|---|---|---|---|---|---|---|
| 1 | 1408 | ACT | GCT | A1408G | 470 | T (Threonine) | A (Alanine) | T470A | 2.24 | -0.11 | Missense | Non-conservative | RNase H |
| 2 | 561 | TTG | TTA | G561A | 187 | L (Leucine) | L(Leucine) | - | | | Silent | | Palm |
| 3 | 284 | CCA | CTA | C284T | 95 | P (Proline) | L(Leucine) | P95L | 0.87 | 0.50 | Missense | Conservative | Palm |
| 4 | 322 | GTA | ATA | G322A | 108 | V (Valine) | I (Isoleucine) | V108M | 1.84 | 0.78 | Missense | Conservative | Palm |
| 5 | 73 | CCA | ACA | C73A | 25 | P (Proline) | T(Threonine) | P25T | 1.18 | 1.04 | Missense | Non-conservative | Finger |
| 6 | 1673, 1674 | AAA | AAG | A1673G, A1674G | 558 | K (Lysine) | K (Lysine) | - | - | - | Silent | | RNase H |
| 6 | 1677 | GTA | GTG | A1677G | 559 | V (Valine) | V (Valine) | - | - | - | Silent | | RNase H |
| 7 | 48 | ATG | ATT | G48T | 16 | M (Methionine) | I (Isoleucine) | M16I | 1.00 | 0.15 | Missense | Conservative | Finger |
| 8 | 1585 | GAA | TAA | G1585T | 529 | E (Glutamic acid) | Stop | E529* | - | - | Nonsense | | RNase H |
| 9 | 1123 | - | - | - | - | - | - | - | - | - | Frameshift | Insertion | |
| 10 | 348 | TTT | TTC | T348C | 116 | F (Phenylalanine) | F(Phenylalanine) | - | - | - | Silent | | Palm |
| 11 | 612 | GAG | GAT | G612T | 204 | E (Glutamic acid) | D (Aspartic acid) | E204D | 1.74 | 0.48 | Missense | Conservative | Palm |
| 12 | 1154 | - | - | - | - | - | - | - | - | - | Frameshift | Deletion | Connection |
| 13 | 220 | TTA | CTA | T220C | 74 | L (Leucine) | L(Leucine) | - | - | - | Silent | | Finger |
| 14 | 1653 | TTA | TTG | A1653G | 551 | L (Leucine) | L(Leucine) | - | - | - | Silent | | RNase H |
| 15A,B | 1661 | GCT | GAT | C1661A | 554 | A (Alanine) | D (Aspartic acid) | A554D | 1.91 | -0.15 | Missense | Non-conservative | RNase H |
| 16 | 413 – 1451 | - | - | - | - | - | - | - | - | - | Frameshift | Deletion | Finger to RNase H |
| 17 | 1 – 1616 | - | - | - | - | - | - | - | - | - | Frameshift | Deletion | Finger to RNase H |
| 18 | 140 – 1631 | - | - | - | - | - | - | - | - | - | Frameshift | Deletion | Finger to RNase H |

| | | | | | | | | | | | | | |
|---|---|---|---|---|---|---|---|---|---|---|---|---|---|
| 19 | 248 | GGA | GTA | G248T | 83 | G (Glycine) | V (Valine) | - | - | - | Silent | | Finger |
| 20 | 104 – 1681 | - | - | - | - | - | - | - | - | - | Frameshift | Deletion | Finger to RNase H |
| 21 | 1258 | CCT | ACT | C1258A | 420 | P (Proline) | T(Threonine) | P420T | 2.02 | 1.68 | Missense | Non-conservative | Connection |
| 22 | 85 – 1616 | - | - | - | - | - | - | - | - | - | Frameshift | Deletion | Finger to RNase H |
| 23 | 717 | TGG | TGT | G717T | 239 | W (Tryptophan) | C (Cysteine) | W239C | 5.79 | 5.20 | Missense | Conservative | Thumb |
| 24 | 345 | TAT | TAC | T345C | 115 | Y (Tyrosine) | Y (Tyrosine) | - | - | - | Silent | | Palm |
| 25 | 683 | CTT | CCT | T683C | 228 | L (Leucine) | P (Proline) | L228P | 1.69 | 1.59 | Missense | Conservative | Thumb |
| 26 | 837 | TTA | TTG | A837G | 279 | L (Leucine) | L(Leucine) | - | - | - | Silent | | Thumb |
| 26 | 969 | AAA | AAG | A969G | 323 | K (Lysine) | K (Lysine) | - | - | - | Silent | | Connection |
| 26 | 975 | TTA | TTG | A975G | 325 | L (Leucine) | L(Leucine) | - | - | - | Silent | | Connection |
| 26 | 976, 978 | ATA | GTG | A976G, A978G | 326 | I (Isoleucine) | V (Valine) | I326V | 0.09 | 0.31 | Missense | Conservative | Connection |
| 26 | 981 | GCA | GCG | A981G | 327 | A (Alanine) | A (Alanine) | - | - | - | Silent | | Connection |
| 26 | 992 | AAG | AGG | A992G | 331 | K (Lysine) | R (Arginine) | K331R | 4.20 | 0.56 | Missense | Conservative | Connection |
| 26 | 1031 | GAG | GGG | A1031G | 344 | E (Glutamic acid) | G (Glycine) | E344G | 0.98 | 0.90 | Missense | Non-conservative | Connection |
| 26 | 1066 | AGA | GGA | A1066G | 356 | R (Arginine) | G (Glycine) | R356G | -0.53 | 0.66 | Missense | Non-conservative | Connection |
| 26 | 1087, 1088 | AAT | GGT | A1087G, A1088G | 363 | N (Asparagine) | A (Alanine) | N363A | 1.85 | 1.22 | Missense | Non-conservative | Connection |
| 26 | 1096 | AAA | GAA | A1096G | 366 | K (Lysine) | E (Glutamic acid) | K366E | 0.30 | 1.40 | Missense | Non-conservative | Connection |
| 26 | 1125 | ATA | ATG | A1125G | 375 | I (Isoleucine) | M (Methionine) | I375M | 1.73 | 1.27 | Missense | Conservative | Connection |
| 27 | 57 – 1625 | - | - | - | - | - | - | - | - | - | Frameshift | Deletion | Finger to RNase H |
| 28 | 285 – 1488 | - | - | - | - | - | - | - | - | - | Frameshift | Deletion | Palm to RNase H |
| 29 | 1073 | AGG | ATG | G1073T | 358 | R (Arginine) | M (Methionine) | R358G | -0.32 | 0.70 | Missense | Non-conservative | Connection |
| 30 | 1606 | GTA | ATA | G1606A | 536 | V (Valine) | I (Isoleucine) | V536I | 2.41 | 0.95 | Missense | Conservative | RNase H |
| 31 | 157 | GAA | TAA | G157T | 53 | E (Glutamic acid) | Stop | E53* | - | - | Nonsense | | Finger |
| 32 | 64 | AAA | GAA | A64G | 22 | K (Lysine) | E (Glutamic acid) | K22E | 0.95 | -0.92 | Missense | Non-conservative | Finger |
| 32 | 237 | GAA | GAG | A237G | 79 | E (Glutamic acid) | E (Glutamic acid) | - | - | - | Silent | | Finger |
| 32 | 267 | GAA | GAG | A267G | 89 | E (Glutamic acid) | E (Glutamic acid) | - | - | - | Silent | | Palm |
| 32 | 308 | AAA | AGA | A308G | 103 | K (Lysine) | R (Arginine) | K103R | -0.09 | 0.11 | Missense | Conservative | Palm |

| | | | | | | | | | | | | | |
|---|---|---|---|---|---|---|---|---|---|---|---|---|---|
| 32 | 406 | AAC | GAC | A406G | 136 | N (Asparagine) | D (Aspartic acid) | N136D | 0.95 | 0.46 | Missense | Non-conservative | Finger |
| 33 | 293 | GCA | GTA | C293T | 98 | A (Alanine) | V (Valine) | A98V | 3.72 | 1.44 | Missense | Conservative | Palm |
| 34 | 1110 | GAG | GAT | G1110T | 370 | E (Glutamic acid) | D (Aspartic acid) | E370D | 1.03 | 1.34 | Missense | Conservative | Connection |
| 35 | 1 – 1579 | - | - | - | - | - | - | - | - | - | Frameshift | Deletion | Finger to RNase H |
| 36 | 861 | - | - | - | - | - | - | - | - | - | Frameshift | Deletion | Thumb |
| 37 | 1140 | ATA | ATG | A1140G | 380 | I (Isoleucine) | M (Methionine) | I380M | 2.52 | -0.16 | Missense | Conservative | Connection |
| 37 | 1221 | CAA | CAG | A1221G | 407 | Q (Glutamine) | Q (Glutamine) | - | - | - | Silent | | Connection |
| 37 | 1252, 1253 | AAT | GGT | A1252G, A1253G | 418 | N (Asparagine) | G (Glycine) | N418G | 1.40 | 1.01 | Missense | Non-conservative | Connection |
| 37 | 1287 | TTA | TTG | A1287G | 429 | L (Leucine) | L(Leucine) | - | - | - | Silent | | RNase H |
| 37 | 1582 | AAG | GAG | A1582G | 528 | K (Lysine) | E (Glutamic acid) | K528E | 3.15 | 5.17 | Missense | Non-conservative | RNase H |
| 38 | 993 | AAG | AAA | G993A | 331 | K (Lysine) | K (Lysine) | - | - | - | Silent | | Connection |
| 39 | 641 | CTT | CCT | T641C | 214 | L (Leucine) | P (Proline) | L214P | 6.94 | 7.07 | Missense | Conservative | Palm |
| 40A,B | 996 | CAG | CAA | G996A | 332 | Q (Glutamine) | Q (Glutamine) | - | - | - | Silent | | Connection |
| 40A,B | 997 | GGG | AGG | G997A | 333 | G (Glycine) | R (Arginine) | G333R | 2.41 | 3.27 | Missense | Non-conservative | Connection |
| 40A,B | 999 | - | - | - | - | - | - | - | - | - | Frameshift | Insertion | Connection |
| 41 | 111 | ATT | ATC | T111C | 37 | I (Isoleucine) | I (Isoleucine) | - | - | - | Silent | | Finger |
| 42 | 182 | TTT | TCT | T182C | 61 | F (Phenylalanine) | S (Serine) | F61S | 1.85 | -0.14 | Missense | Non-conservative | Finger |
| 42 | 1553 | GTC | GCC | T1553C | 518 | V (Valine) | A (Alanine) | V518A | 2.94 | 2.35 | Missense | Conservative | RNase H |

**Table S4 Codon mutated HIV-1 RT p66 nucleotide and amino acid mutations.** Free energy values $\Delta\Delta G_{Mt}$ (kcal / mol) were rounded to 2 decimal places.

| Variant | Nucleotide position | Initial codon | Mutated codon | Nucleotide Mutation | Amino Acid Position | Initial Amino Acid | Mutated Amino Acid | Amino Acid Mutation | Rosetta Cartesian_ddg ΔΔGMt (kcal / mol) | FoldX BuildPDB ΔΔGMt (kcal / mol) | Type of mutation | | Domain |
|---|---|---|---|---|---|---|---|---|---|---|---|---|---|
| 1 | 1477 | GCT | TCT | G1477T | 493 | A (Alanine) | S (Serine) | A493S | 1.66 | 0.14 | Missense | Non-conservative | RNase H |
| 2 | 199 | AAA | GAA | A199G | 67 | K (Lysine) | E (Glutamic Acid) | K67E | 0.36 | -0.17 | Missense | Non-conservative | Finger |
| 3 | 1384 | ACA | TCA | A1384T | 462 | T (Threonine) | S (Serine) | T462S | 1.30 | 0.50 | Missense | Conservative | RNase H |
| 4A,B | 636 | TTT | TTC | T636C | 212 | F (Phenylalanine) | F (Phenylalanine) | - | - | - | Silent | | Palm |
| 5A,B | 898 | CAT | AAT | C898A | 300 | H (Histidine) | N (Asparagine) | H300N | 2.18 | -0.01 | Missense | Non-conservative | Thumb |
| 6 | 1198 | GAA | AAA | G1198A | 400 | E (Glutamic Acid) | K (Lysine) | E400K | 3.00 | -0.20 | Missense | Non-conservative | Connection |
| 7 | 657 | TTA | TTG | A657G | 219 | L (Leucine) | L (Leucine) | - | - | - | Silent | | Palm |
| 7 | 792 | TTA | TTG | A792G | 264 | L (Leucine) | L (Leucine) | - | - | - | Silent | | Thumb |
| 7 | 805 | AGA | GGA | A805G | 269 | R (Arginine) | G (Glycine) | R269G | 2.16 | 0.67 | Missense | Non-conservative | Thumb |
| 7 | 828 | GAA | GAG | A828G | 276 | E (Glutamic Acid) | E (Glutamic Acid) | - | - | - | Silent | | Thumb |
| 7 | 840 | TTA | TTG | A840G | 280 | L (Leucine) | L (Leucine) | - | - | - | Silent | | Thumb |
| 7 | 841 | ACA | GCA | A841G | 281 | T (Threonine) | A (Alanine) | T281A | 1.86 | 0.06 | Missense | Non-conservative | Thumb |
| 7 | 922, 924 | AAA | GAG | A922G, A924G | 308 | K (Lysine) | E (Glutamic Acid) | K308E | 3.22 | 0.74 | Missense | Non-conservative | Connection |
| 7 | 930 | TTA | TTG | A930G | 310 | L (Leucine) | L (Leucine) | - | - | - | Silent | | Connection |
| 7 | 1050 | GTA | GTG | A1050G | 350 | V (Valine) | V (Valine) | - | - | - | Silent | | Connection |
| 7 | 1051 | AAA | GAA | A1051G | 351 | K (Lysine) | E (Glutamic Acid) | K351E | 0.30 | 1.40 | Missense | Non-conservative | Connection |
| 8A,B | 1165 | GAA | TAA | G1165T | 389 | E (Glutamic Acid) | Stop | E389* | - | - | Nonsense | | Connection |
| 9 | 1501 | GAA | AAA | G1501A | 501 | E (Glutamic Acid) | K (Lysine) | E501K | 0.41 | 0.65 | Missense | Non-conservative | RNase H |

| | | | | | | | | | | | | | |
|---|---|---|---|---|---|---|---|---|---|---|---|---|---|
| 10 | 109 | CCA | ACA | C109A | 37 | P (Proline) | T (Threonine) | P37T | 2.44 | 0.70 | Missense | Non-conservative | Finger |
| 11 | 809 | GGA | GAA | G809A | 270 | G (Glycine) | E (Glutamic Acid) | G270E | 6.00 | 3.31 | Missense | Non-conservative | Thumb |
| 12 | 1157 | TGG | TTG | G1157T | 386 | W (Tryptophan) | L (Leucine) | W386L | 2.79 | 2.86 | Missense | Conservative | Connection |
| 13 | 822 | TTA | TTT | A822T | 274 | L (Leucine) | F (Phenylalanine) | L274F | 2.11 | 0.44 | Missense | Conservative | Thumb |
| 14 | 1214 | CCA | CAA | C1214A | 405 | P (Proline) | Q (Glutamine) | P405Q | 2.96 | 1.37 | Missense | Non-conservative | Connection |
| 15 | 89 | GGA | GAA | G89A | 30 | G (Glycine) | E (Glutamic Acid) | G30E | 1.06 | 4.80 | Missense | Non-conservative | Finger |
| 16 | 1180 | ACA | GCA | A1180G | 394 | T (Threonine) | A (Alanine) | T394A | 2.00 | 0.45 | Missense | Non-conservative | Connection |
| 17 | 1150 | GAA | TAA | G1150T | 384 | E (Glutamic Acid) | Stop | E384* | - | - | Nonsense | | Connection |
| 18A,B,C | 1428 | TTA | TTG | A1428G | 476 | L (Leucine) | L (Leucine) | - | - | - | Silent | | RNase H |
| 18A,B,C | 1444 | ACA | GCA | A1444G | 482 | T (Threonine) | A (Alanine) | T482A | 2.66 | 1.71 | Missense | Non-conservative | RNase H |
| 18A,B,C | 1510 | AAT | GAT | A1510G | 504 | N (Asparagine) | D (Aspartic Acid) | N504D | 2.49 | 1.30 | Missense | Non-conservative | RNase H |
| 18A,B,C | 1519 | ATT | GTT | A1519G | 507 | I (Isoleucine) | V (Valine) | I507V | 2.75 | 1.20 | Missense | Conservative | RNase H |
| 18A,B,C | 1530 | TTA | TTG | A1530G | 510 | L (Leucine) | L (Leucine) | - | - | - | Silent | | RNase H |
| 18A,B,C | 1573 | AAA | GAA | A1573G | 525 | K (Lysine) | E (Glutamic Acid) | K525E | 1.55 | 0.58 | Missense | Non-conservative | RNase H |
| 18A,B,C | 1611 | GTA | GTG | A1611G | 537 | V (Valine) | V (Valine) | - | - | - | Silent | | RNase H |
| 18A,B,C | 1621 | ATT | GTT | A1621G | 541 | I (Isoleucine) | V (Valine) | I541V | 1.17 | 1.46 | Missense | Conservative | RNase H |
| 18A,B,C | 1627 | AAA | GAA | A1627G | 543 | K (Lysine) | E (Glutamic Acid) | K543E | -0.99 | 0.17 | Missense | Non-conservative | RNase H |
| 18A,B,C | 1635 | TTA | TTG | A1635G | 545 | L (Leucine) | L (Leucine) | - | - | - | Silent | | RNase H |
| 19 | 1444 - 1473 | - | - | - | - | - | - | - | - | - | Frameshift | Deletion | RNase H |
| 20 | 989 | CCA | CAA | C989A | 330 | P (Proline) | Q (Glutamine) | P330Q | 0.76 | 1.20 | Missense | Non-conservative | Connection |

| | | | | | | | | | | | | | |
|---|---|---|---|---|---|---|---|---|---|---|---|---|---|
| 21 | 994 | AAA | GAA | A994G | 332 | K (Lysine) | E (Glutamic Acid) | K332E | 1.44 | 0.85 | Missense | Non-conservative | Connection |
| 21 | 1016 | TAT | TGT | A1016G | 339 | Y (Tyrosine) | C (Cysteine) | Y339C | 3.58 | 2.75 | Missense | Non-conservative | Connection |
| 21 | 1050 | GTA | GTG | A1050G | 350 | V (Valine) | V (Valine) | - | - | - | Silent | | Connection |
| 21 | 1053 | AAA | AAG | A1053G | 351 | K (Lysine) | K (Lysine) | - | - | - | Silent | | Connection |
| 21 | 1077 | AAA | AAG | A1077G | 359 | K (Lysine) | K (Lysine) | - | - | - | Silent | | Connection |
| 21 | 1099 | ATT | GTT | A1099G | 367 | I (Isoleucine) | V (Valine) | I367V | 1.98 | 0.97 | Missense | Conservative | Connection |
| 21 | 1123 | AAA | GAA | A1123G | 375 | K (Lysine) | E (Glutamic Acid) | K375E | -0.24 | 3.15 | Missense | Non-conservative | Connection |
| 21 | 1139, 1140 | AAA | AGG | A1139G, A1140G | 380 | K (Lysine) | R (Arginine) | K380R | -0.04 | 0.53 | Missense | Conservative | Connection |
| 21 | 1169 | TAT | TGT | A1169G | 390 | Y (Tyrosine) | C (Cysteine) | Y390C | 2.52 | 2.50 | Missense | Non-conservative | Connection |
| 21 | 1180, 1182 | ACA | GCG | A1180G, A1182G | 394 | T (Threonine) | A (Alanine) | T394A | 2.00 | 0.45 | Missense | Non-conservative | Connection |
| 21 | 1221 | TTA | TTG | A1221G | 407 | L (Leucine) | L (Leucine) | - | - | - | Silent | | Connection |
| 21 | 1242 | TTA | TTG | A1242G | 414 | L (Leucine) | L (Leucine) | - | - | - | Silent | | RNase H |
| 21 | 1277 | TAT | TGT | A1277G | 426 | Y (Tyrosine) | C (Cysteine) | Y426C | 4.02 | 2.42 | Missense | Non-conservative | RNase H |
| 21 | 1384 | ACA | GCA | A1384G | 462 | T (Threonine) | A (Alanine) | T462A | -0.17 | -0.95 | Missense | Non-conservative | RNase H |
| 21 | 1392 | TTA | TTG | A1392G | 464 | L (Leucine) | L (Leucine) | - | - | - | Silent | | RNase H |
| 21 | 1399 | ATT | GTT | A1399G | 467 | I (Isoleucine) | V (Valine) | I467V | 1.64 | 0.94 | Missense | Conservative | RNase H |
| 21 | 1407 | TTA | TTG | A1407G | 469 | L (Leucine) | L (Leucine) | - | - | - | Silent | | RNase H |
| 21 | 1434 | GTA | GTG | A1434G | 478 | V (Valine) | V (Valine) | - | - | - | Silent | | RNase H |
| 21 | 1435 | AAT | GAT | A1435G | 479 | N (Asparagine) | D (Aspartic Acid) | N479D | 1.32 | 2.63 | Missense | Non-conservative | RNase H |
| 21 | 1464 | TTA | TTG | A1464G | 488 | L (Leucine) | L (Leucine) | - | - | - | Silent | | RNase H |

| | | | | | | | | | | | | | |
|---|---|---|---|---|---|---|---|---|---|---|---|---|---|
| 21 | 1530 | TTA | TTG | A1530G | 510 | L (Leucine) | L (Leucine) | - | - | - | Silent | | RNase H |
| 21 | 1531 | ATT | GTT | A1531G | 511 | I (Isoleucine) | V (Valine) | I511V | 1.50 | 0.81 | Missense | Conservative | RNase H |
| 22 | 88 | GGA | TGA | G88T | 30 | G (Glycine) | Stop | G30* | - | - | Nonsense | | Finger |
| 23 | 1474 | CAA | AAA | C1474A | 492 | Q (Glutamine) | K (Lysine) | Q492K | 0.49 | -0.65 | Missense | Non-conservative | RNase H |
| 24 | 153 | - | - | - | - | - | - | - | - | - | Frameshift | Insertion | |
| 25 | 1232 | TGG | TTG | G1232T | 411 | W (Tryptophan) | L (Leucine) | W411L | 0.60 | -0.15 | Missense | Conservative | Connection |
| 26 | 1063 | GAA | TAA | G1063T | 355 | E (Glutamic Acid) | Stop | E355* | - | - | Nonsense | | Connection |
| 27 | 800 | CTT | CCT | T800C | 267 | L (Leucine) | P (Proline) | L267P | 8.25 | 5.17 | Missense | Conservative | Thumb |
| 28 | 568 | TTA | CTA | T568C | 190 | L (Leucine) | L (Leucine) | - | - | - | Silent | | Palm |
| 29 | 1396 | GCA | TCA | G1396T | 466 | A (Alanine) | S (Serine) | A466S | 2.66 | 0.98 | Missense | Non-conservative | RNase H |
| 30 | 723 | GAT | GAC | T723C | 241 | D (Aspartic Acid) | D (Aspartic Acid) | - | - | - | Silent | | Thumb |
| 31 | 682 | CCA | TCA | C682T | 228 | P (Proline) | S (Serine) | P228S | 2.32 | 1.56 | Missense | Non-conservative | Thumb |
| 32 | 21 | AAA | AAG | A21G | 7 | K (Lysine) | K (Lysine) | - | - | - | Silent | | Finger |
| 32 | 33 | TTA | TTG | A33G | 11 | L (Leucine) | L (Leucine) | - | - | - | Silent | | Finger |
| 32 | 39 | GAA | GAG | A39G | 13 | E (Glutamic Acid) | E (Glutamic Acid) | - | - | - | Silent | | Finger |
| 32 | 44 | AAA | AGA | A44G | 15 | K (Lysine) | R (Arginine) | K15R | 0.84 | -0.01 | Missense | Conservative | Finger |
| 32 | 57 | TTA | TTG | A57G | 19 | L (Leucine) | L (Leucine) | - | - | - | Silent | | Finger |
| 32 | 70 | ACA | GCA | A70G | 24 | T (Threonine) | A (Alanine) | T24A | 1.51 | -0.13 | Missense | Non-conservative | Finger |
| 32 | 84 | AAA | AAG | A84G | 28 | K (Lysine) | K (Lysine) | - | - | - | Silent | | Finger |
| 32 | 100, 102 | AAA | GAG | A100G, A102G | 34 | K (Lysine) | E (Glutamic Acid) | K34E | 3.11 | 1.24 | Missense | Non-conservative | Finger |

| | | | | | | | | | | | | | |
|---|---|---|---|---|---|---|---|---|---|---|---|---|---|
| 32 | 142 | ATT | GTT | A142G | 48 | I (Isoleucine) | V (Valine) | I48V | 1.75 | 0.68 | Missense | Conservative | Finger |
| 32 | 146 | AAG | AGG | A146G | 49 | K (Lysine) | R (Arginine) | K49R | -2.23 | 0.22 | Missense | Conservative | Finger |
| 32 | 151, 153 | AAA | GAG | A151G, A153G | 51 | K (Lysine) | E (Glutamic Acid) | K51E | 3.38 | -0.08 | Missense | Non-conservative | Finger |
| 32 | 165 | AAA | AAG | A165G | 55 | K (Lysine) | K (Lysine) | - | - | - | Silent | | Finger |
| 32 | 177 | TTA | TTG | A177G | 59 | L (Leucine) | L (Leucine) | - | - | - | Silent | | Finger |
| 32 | 364 | AAT | GAT | A364G | 122 | N (Asparagine) | D (Aspartic Acid) | N122D | -0.47 | 0.52 | Missense | Non-conservative | Finger |
| 33 | 225 | GTA | GTG | A225G | 75 | V (Valine) | V (Valine) | - | - | - | Silent | - | Palm |
| 33 | 784 | CGT | AGT | C784A | 262 | R (Arginine) | S (Serine) | R262S | 1.20 | 2.08 | Missense | Non-conservative | Thumb |
| 34 | 1010 | GGA | GTA | G1010T | 337 | G (Glycine) | V (Valine) | G337V | 2.49 | 14.36 | Missense | Conservative | Connection |
| 35 | 377 – 401 | - | - | - | - | - | - | - | - | - | Frameshift | Deletion | Finger |
| 36 | 1127 | CTT | CCT | T1127C | 376 | L (Leucine) | P (Proline) | L376P | 5.83 | 8.53 | Missense | Conservative | Connection |
| 37A,B | 460 | GAA | AAA | G460A | 154 | E (Glutamic Acid) | K (Lysine) | E154K | 1.54 | -0.19 | Missense | Non-conservative | Palm |
| 38 | 1429 | GAA | AAA | G1429A | 477 | E (Glutamic Acid) | K (Lysine) | E477K | 1.13 | -0.75 | Missense | Non-conservative | RNase H |
| 39 | 1573 | AAA | GAA | A1573G | 525 | K (Lysine) | E (Glutamic Acid) | K525E | 1.55 | 0.58 | Missense | Non-conservative | RNase H |
| 39 | 1579 | ATT | GTT | A1579G | 527 | I (Isoleucine) | V (Valine) | I527V | -0.26 | 0.63 | Missense | Conservative | RNase H |
| 39 | 1588 | AAT | GAT | A1588G | 530 | N (Asparagine) | D (Aspartic Acid) | N530D | 1.33 | 1.47 | Missense | Non-conservative | RNase H |
| 39 | 1596 | CAA | CAG | A1596G | 532 | Q (Glutamine) | Q (Glutamine) | - | - | - | Silent | | RNase H |
| 39 | 1599 | GTA | GTG | A1599G | 533 | V (Valine) | V (Valine) | - | - | - | Silent | | RNase H |
| 39 | 1604 | AAA | AGA | A1604G | 535 | K (Lysine) | R (Arginine) | K535R | 1.04 | 0.05 | Missense | Conservative | RNase H |
| 39 | 1624 | AGA | GGA | A1624G | 542 | R (Arginine) | G (Glycine) | R542G | 0.75 | 0.80 | Missense | Non-conservative | RNase H |

| | | | | | | | | | | | | | |
|---|---|---|---|---|---|---|---|---|---|---|---|---|---|
| 39 | 1632 | GTA | GTG | A1632G | 544 | V (Valine) | V (Valine) | - | - | - | Silent | | RNase H |
| 39 | 1635 | TTA | TTG | A1635G | 545 | L (Leucine) | L (Leucine) | - | - | - | Silent | | RNase H |
| 40 | 1069 | GTT | ATT | G1069A | 357 | V (Valine) | I (Isoleucine) | V357I | 1.18 | -0.95 | Missense | Conservative | Connection |
| 41 | 634 | TTT | ATT | T634A | 212 | F (Phenylalanine) | I (Isoleucine) | F212I | 5.04 | 3.25 | Missense | Conservative | Palm |
| 42 | 1242 | TTA | TTG | A1242G | 414 | L (Leucine) | L (Leucine) | - | - | - | Silent | | RNase H |
| 42 | 1248 | AAA | AAG | A1248G | 416 | K (Lysine) | K (Lysine) | - | - | - | Silent | | RNase H |
| 42 | 1260 | GTA | GTG | A1260G | 420 | V (Valine) | V (Valine) | - | - | - | Silent | | RNase H |
| 42 | 1272 | ACA | ACG | A1272G | 424 | T (Threonine) | T (Threonine) | - | - | - | Silent | | RNase H |
| 42 | 1277 | TAT | TGT | A1277G | 426 | Y (Tyrosine) | C (Cysteine) | Y426C | 4.02 | 2.42 | Missense | Non-conservative | RNase H |
| 42 | 1302 | GAA | GAG | A1302G | 434 | E (Glutamic Acid) | E (Glutamic Acid) | - | - | - | Silent | | RNase H |
| 42 | 1307 | AAA | AGA | A1307G | 436 | K (Lysine) | R (Arginine) | K436R | 0.06 | 0.27 | Missense | Conservative | RNase H |
| 42 | 1342 | AGA | GGA | A1342G | 448 | R (Arginine) | G (Glycine) | R448G | 3.91 | 2.31 | Missense | Non-conservative | RNase H |
| 42 | 1349, 1350 | AAA | AGG | A1349G, A1350G | 450 | K (Lysine) | R (Arginine) | K450R | 1.75 | 0.37 | Missense | Conservative | RNase H |
| 42 | 1353 | GTA | GTG | A1353G | 451 | V (Valine) | V (Valine) | - | - | - | Silent | | RNase H |
| 42 | 1381, 1382, 1383 | AAA | GGG | A1381G, A1382G, A1383G | 461 | K (Lysine) | G (Glycine) | K461G | 2.08 | 1.55 | Missense | Non-conservative | RNase H |
| 42 | 1399 | ATT | GTT | A1399G | 467 | I (Isoleucine) | V (Valine) | I467V | 1.64 | 0.94 | Missense | Conservative | RNase H |
| 42 | 1428 | TTA | TTG | A1428G | 476 | L (Leucine) | L (Leucine) | - | - | - | Silent | | RNase H |
| 42 | 1444 | ACA | GCA | A1444G | 482 | T (Threonine) | A (Alanine) | T482A | 2.66 | 1.71 | Missense | Non-conservative | RNase H |
| 42 | 1450 | AGT | GGT | A1450G | 484 | S (Serine) | G (Glycine) | S484G | 3.32 | 0.86 | Missense | Non-conservative | RNase H |
| 42 | 1464 | TTA | TTG | A1464G | 488 | L (Leucine) | L (Leucine) | - | - | - | Silent | | RNase H |

| | | | | | | | | | | | | | |
|---|---|---|---|---|---|---|---|---|---|---|---|---|---|
| 42 | 1471 | ATT | GTT | A1471G | 491 | I (Isoleucine) | V (Valine) | I491V | 2.31 | 1.14 | Missense | Conservative | RNase H |
| 42 | 1476 | CAA | CAG | A1476G | 492 | Q (Glutamine) | Q (Glutamine) | - | - | - | Silent | | RNase H |
| 42 | 1482 | CAA | CAG | A1482G | 494 | Q (Glutamine) | Q (Glutamine) | - | - | - | Silent | | RNase H |
| 42 | 1510 | AAT | GAT | A1510G | 504 | N (Asparagine) | D (Aspartic Acid) | N504D | 2.49 | 1.30 | Missense | Non-conservative | RNase H |
| 42 | 1534, 1535 | AAG | GGG | A1534G, A1535G | 512 | K (Lysine) | G (Glycine) | K512G | 0.74 | 0.21 | Missense | Non-conservative | RNase H |
| 42 | 1542 | GAA | GAG | A1542G | 514 | E (Glutamic Acid) | E (Glutamic Acid) | - | - | - | Silent | | RNase H |
| 42 | 1543 | AAA | GAA | A1543G | 515 | K (Lysine) | E (Glutamic Acid) | K515E | 3.35 | 2.49 | Missense | Non-conservative | RNase H |
| 42 | 1554 | TTA | TTG | A1554G | 518 | L (Leucine) | L (Leucine) | - | - | - | Silent | | RNase H |
| 42 | 1573, 1574 | AAA | GGA | A1573G, A1574G | 525 | K (Lysine) | G (Glycine) | K525G | 1.91 | 1.20 | Missense | Non-conservative | RNase H |
| 42 | 1588 | AAT | GAT | A1588G | 530 | N (Asparagine) | D (Aspartic Acid) | N530D | 1.33 | 1.47 | Missense | Non-conservative | RNase H |
| 42 | 1596 | CAA | CAG | A1596G | 532 | Q (Glutamine) | Q (Glutamine) | - | - | - | Silent | | RNase H |
| 42 | 1599 | GTA | GTG | A1599G | 533 | V (Valine) | V (Valine) | - | - | - | Silent | | RNase H |
| 42 | 1612 | AGT | GGT | A1612G | 538 | S (Serine) | G (Glycine) | S538G | 0.78 | 0.23 | Missense | Non-conservative | RNase H |
| 42 | 1627 | AAA | GAA | A1627G | 543 | K (Lysine) | E (Glutamic Acid) | K543E | -0.99 | 0.17 | Missense | Non-conservative | RNase H |
| 42 | 1635 | TTA | TTG | A1635G | 545 | L (Leucine) | L (Leucine) | - | - | - | Silent | | RNase H |
| 43 | 829 | GTA | ATA | G829A | 277 | V (Valine) | I (Isoleucine) | V277I | -0.02 | -0.31 | Missense | Conservative | Thumb |
| 44 | 1073 | - | - | - | - | - | - | - | - | - | Frameshift | Deletion | Connection |
| 45 | 573 | CGT | CGC | T573C | 191 | R (Arginine) | R (Arginine) | - | - | - | Silent | | Palm |
| 46 | 139 | GCT | ACT | G139A | 47 | A (Alanine) | T (Threonine) | A47T | 1.73 | 0.83 | Missense | Non-conservative | Finger |
| 47A,B,C | 981 | TAT | TAC | T981C | 327 | Y (Tyrosine) | Y (Tyrosine) | - | - | - | Silent | | Connection |

| | | | | | | | | | | | | | |
|---|---|---|---|---|---|---|---|---|---|---|---|---|---|
| 48 | 1030 | GGT | TGT | G1030T | 344 | G (Glycine) | C (Cysteine) | G344C | 1.51 | 3.74 | Missense | Conservative | Connection |
| 48 | 1267 | GAA | TAA | G1267T | 423 | E (Glutamic Acid) | Stop | E423* | - | - | Nonsense | | RNase H |
| 49 | 889 | GAA | AAA | G889A | 297 | E (Glutamic Acid) | K (Lysine) | E297K | 0.34 | -0.08 | Missense | Non-conservative | Thumb |
| 50 | 1002 | TTA | TTG | A1002G | 334 | L (Leucine) | L (Leucine) | - | - | - | Silent | | Connection |
| 50 | 1005 | AAA | AAG | A1005G | 335 | K (Lysine) | K (Lysine) | - | - | - | Silent | | Connection |
| 50 | 1006 | ACA | GCA | A1006G | 336 | T (Threonine) | A (Alanine) | T336A | -0.18 | 1.50 | Missense | Non-conservative | Thumb |
| 50 | 1013, 1014 | AAA | AGG | A1013G, A1014G | 338 | K (Lysine) | R (Arginine) | K338R | 1.88 | 0.45 | Missense | Conservative | Thumb |
| 50 | 1024 | ATG | GTG | A1024G | 342 | M (Methionine) | V (Valine) | M342V | 2.15 | 1.04 | Missense | Conservative | Thumb |
| 50 | 1059 | TTA | TTG | A1059G | 353 | L (Leucine) | L (Leucine) | - | - | - | Silent | | Connection |
| 50 | 1247, 1248 | AAA | AGG | A1247G, A1248G | 416 | K (Lysine) | R (Arginine) | K416R | 0.54 | 0.05 | Missense | Conservative | RNase H |
| 50 | 121428 | TTA | TTG | A1428G | 476 | L (Leucine) | L (Leucine) | - | - | - | Silent | - | RNase H |
| 50 | 1431 | GAA | GAG | A1431G | 477 | E (Glutamic Acid) | E (Glutamic Acid) | - | - | - | Silent | - | RNase H |
| 50 | 1464 | TTA | TTG | A1464G | 488 | L (Leucine) | L (Leucine) | - | - | - | Silent | - | RNase H |
| 50 | 1471 | ATT | GTT | A1471G | 491 | I (Isoleucine) | V (Valine) | I491V | 2.31 | 1.14 | Missense | Conservative | RNase H |
| 50 | 1476 | CAA | CAG | A1476G | 492 | Q (Glutamine) | Q (Glutamine) | - | - | - | Silent | - | RNase H |
| 51 | 1139, 1140 | AAA | AGG | A1139G, A1140G | 380 | K (Lysine) | R (Arginine) | K380R | -0.04 | 0.53 | Missense | Conservative | Connection |
| 52 | 247 | GCA | TCA | G247T | 83 | A (Alanine) | S (Serine) | A83S | 3.19 | 1.26 | Missense | Non-conservative | Palm |
| 53 | 892 | CCA | TCA | C892T | 298 | P (Proline) | S (Serine) | P298S | -0.38 | 0.88 | Missense | Non-conservative | Thumb |
| 54 | 1 – 233 | - | - | - | - | - | - | - | - | - | Frameshift | Deletion | Finger to Palm |
| 55 | 1 – 959 | - | - | - | - | - | - | - | - | - | Frameshift | Deletion | Finger to Connection |

| | | | | | | | | | | | | | |
|---|---|---|---|---|---|---|---|---|---|---|---|---|---|
| 56 | 1464 | TTA | TTG | A1464G | 488 | L (Leucine) | L (Leucine) | - | - | - | Silent | - | RNase H |
| 57 | 893 | CCA | CTA | C893T | 298 | P (Proline) | L (Leucine) | P298L | 1.38 | 0.53 | Missense | Conservative | Thumb |
| 58 | 185 | TTT | TCT | T185C | 62 | F (Phenylalanine) | S (Serine) | F62S | 3.73 | 3.96 | Missense | Non-conservative | Finger |
| 59 | 1143 | GAA | GAG | A1143G | 381 | E (Glutamic Acid) | E (Glutamic Acid) | - | - | - | Silent | - | Connection |
| 60 | 1447 | GAT | TAT | G1447T | 483 | D (Aspartic Acid) | Y (Tyrosine) | D483Y | -1.02 | -1.91 | Missense | Non-conservative | RNase H |
| 61A,B | 1221 | TTA | TTG | A1221G | 407 | L (Leucine) | L (Leucine) | - | - | - | Silent | - | Connection |
| 61A,B | 1225 | AAA | GAA | A1225G | 409 | K (Lysine) | E (Glutamic Acid) | K409E | 2.45 | 0.09 | Missense | Non-conservative | Connection |
| 61A,B | 1260 | GTA | GTG | A1260G | 420 | V (Valine) | V (Valine) | - | - | - | Silent | - | RNase H |
| 61A,B | 1263 | GGA | GGG | A1263G | 421 | G (Glycine) | G (Glycine) | - | - | - | Silent | - | RNase H |
| 61A,B | 1297 | AGA | GGA | A1297G | 433 | R (Arginine) | G (Glycine) | R433G | -0.78 | 0.18 | Missense | Non-conservative | RNase H |
| 61A,B | 1308 | AAA | AAG | A1308G | 436 | K (Lysine) | K (Lysine) | - | - | - | Silent | | RNase H |
| 62 | 1513 | CAA | AAA | C1513A | 505 | Q (Glutamine) | K (Lysine) | Q505K | 0.04 | 0.07 | Missense | Non-conservative | RNase H |
| 63 | 970 | TAT | GAT | T970G | 324 | Y (Tyrosine) | D (Aspartic Acid) | Y324D | 3.80 | 6.55 | Missense | Non-conservative | Connection |
| 64 | 167 | TGG | TTG | G167T | 56 | W (Tryptophan) | L (Leucine) | W56L | 2.82 | 1.21 | Missense | Conservative | Finger |
| 65 | 821 | TTA | TCA | T821C | 274 | L (Leucine) | S (Serine) | L274S | 0.34 | 1.65 | Missense | Non-conservative | Thumb |
| 66 | 1172 | TGG | TTG | G1172T | 391 | W (Tryptophan) | L (Leucine) | W391L | 0.77 | -0.15 | Missense | Conservative | Connection |
| 67 | 114 | GAA | GAG | A114G | 38 | E (Glutamic Acid) | E (Glutamic Acid) | - | - | - | Silent | - | Finger |
| 67 | 135 | GTA | GTG | A135G | 45 | V (Valine) | V (Valine) | - | - | - | Silent | - | Finger |
| 67 | 145, 146 | AAG | GGG | A145G, A146G | 49 | K (Lysine) | G (Glycine) | K49G | 4.06 | 1.06 | Missense | Non-conservative | Finger |
| 68 | 895 | GTA | ATA | G895A | 299 | V (Valine) | I (Isoleucine) | V299I | 0.03 | -0.03 | Missense | Conservative | Thumb |

| | | | | | | | | | | | | | |
|---|---|---|---|---|---|---|---|---|---|---|---|---|---|
| 69 | 371 | ACA | AAA | C371A | 124 | T (Threonine) | K (Lysine) | T124K | 1.70 | -0.85 | Missense | Non-conservative | Finger |
| 70 | 772 | GGT | AGT | G772A | 258 | G (Glycine) | S (Serine) | G258S | 2.20 | 3.78 | Missense | Non-conservative | Thumb |
| 71 | 1247, 1248 | AAA | AGG | A1247G, A1248G | 416 | K (Lysine) | R (Arginine) | K416R | 0.54 | 0.05 | Missense | Conservative | RNase H |
| 71 | 1325 | TAT | TGT | A1325G | 442 | Y (Tyrosine) | C (Cysteine) | Y442C | 3.52 | 3.54 | Missense | Non-conservative | RNase H |
| 71 | 1330 | ACA | GCA | A1330G | 444 | T (Threonine) | A (Alanine) | T444A | 2.34 | 3.61 | Missense | Non-conservative | RNase H |
| 71 | 1403 | TAT | TGT | A1403G | 468 | Y (Tyrosine) | C (Cysteine) | Y468C | 3.25 | 1.94 | Missense | Non-conservative | RNase H |
| 72 | 137 – 168 | - | - | - | - | - | - | - | - | - | Frameshift | Deletion | Finger |
| 73 | 436 | CAA | TAA | C436T | 146 | Q (Glutamine) | Stop | Q146* | - | - | Nonsense | - | Palm |
| 74 | 1016 | TAT | TGT | A1016G | 339 | Y (Tyrosine) | C (Cysteine) | Y339C | 3.58 | 2.75 | Missense | Non-conservative | Connection |
| 74 | 1065 | GAA | GAG | A1065G | 355 | E (Glutamic Acid) | E (Glutamic Acid) | - | - | - | Silent | - | Connection |
| 75 | 587 | AGA | ATA | G587T | 196 | R (Arginine) | I (Isoleucine) | R196I | 0.35 | 0.03 | Missense | Non-conservative | Palm |
| 76 | 998 | AAT | AGT | A998G | 333 | N (Asparagine) | S (Serine) | N333S | 2.15 | 1.03 | Missense | Conservative | Connection |
| 77 | 1444 | ACA | GCA | A1444G | 482 | T (Threonine) | A (Alanine) | T482A | 2.66 | 1.71 | Missense | Non-conservative | RNase H |
| 78 | 666 | GAT | GAC | T666C | 222 | N (Asparagine) | N (Asparagine) | - | - | - | Silent | - | Palm |
| 79 | 882 | ATT | ATC | T882C | 294 | I (Isoleucine) | I (Isoleucine) | - | - | - | Silent | - | Thumb |
| 80 | 167 | TGG | TAG | G167A | 56 | W (Tryptophan) | Stop | W56* | - | - | Nonsense | - | Finger |
| 81 | 710 | TGG | TTG | G710T | 237 | W (Tryptophan) | L (Leucine) | W237L | 4.80 | 0.48 | Missense | Conservative | Thumb |
| 82 | 876 | CGT | CGC | T876C | 292 | R (Arginine) | R (Arginine) | - | - | - | Silent | - | Thumb |
| 83 | 1219 | TTA | CTA | T1219C | 407 | L (Leucine) | L (Leucine) | - | - | - | Silent | - | Connection |
| 84A,B | 694 - 717 | - | - | - | - | - | - | - | - | - | - | Deletion | Thumb |

**Table S5 Clinical, drug resistant and mutations with reported functions of HIV-1 Gag, Protease and RT p66.**

| Mutation | Domain | Implication / Reported Function | Type | Reference |
|---|---|---|---|---|
| **HIV-1 Gag** | | | | |
| E17K | Matrix | CTL immune evasion resistance.<br>Rare and transient mutation. | Clinical Isolate; In Vitro | (1, 2) |
| | Matrix | E17K was acquired when selecting with novel PI GRL-0519, along with V84A, G61E and D152N | In Vitro | (3) |
| E42K | Matrix | Compensatory mutations (E42K and P10L) responsible for enhanced infectivity, to overcome deletion in stem-loop 1 | In vivo | (4) |
| G192R | Capsid | G192W (denoted as G60W) increased the number of viral particles at N-terminal domain of capsid protein | In vitro | (5) |
| K202R | Capsid | Part of PF74 (capsid inhibitor) binding site with Q67H and T107N.<br>Conferred low-level resistance to capsid inhibitor (PF74), impaired HIV-1 infectivity by 90% and reduced PF74 binding to HIV-1 particles | In vitro | (6) |
| R214G | Capsid | Alanine scanning revealed 3-fold decrease in infectivity | In vitro | (7) |
| P222L | Capsid | Cyclophilin A (CyPA) binding site (along with G221), with mutant P222A found to disrupt binding to CyPA. | In vitro | (8) |
| I223V | Capsid | CTL immune evasion resistance.<br>Known compensatory mutation for T242N escape mutation. | In vitro | (9, 10) |
| S241I | Capsid | Transiently observed variant (TiTLQEQIGW) with no reported function | Clinical Isolate | (11) |
| E245K | Capsid | E245D found to have diminished IFN-γ response | In vivo | (12) |
| N271S | Capsid | CTL immune evasion resistance.<br>Known rare and transient mutation. | In vitro | (13) |
| K290R | Capsid | Did not respond to inositol hexakisphosphate (IP6) and s-CANC, consistent with high degree of lysine conservation | In vitro | (14) |
| A431D | Nucleocapsid | Amino acid position 431 found to be influenced by positive selection.<br>A431V found in patient after PI treatment. | Clinical Isolate | (15) |
| | | Reduced susceptibility to ritonavir by 3.8-fold | In vitro | (16) |
| P453T | P6 | Amino acid 453 found to be influenced by positive selection.<br>P453L found in patient after PI treatment. | Clinical Isolate | (15) |
| | | L449F/P453T were selected after high-pressure passage with protease inhibitor GW640385 | In vitro | (17) |
| T470A | P6 | Associated with reduced replication capacity, with polymorphism increasing over course of epidemic in Japan | Clinical Isolate | (18) |
| | | Indicated as an escape variant, peptide titration using PBMCs of HLA-Cw*08 patient demonstrated that peptide containing T470A was more weakly recognised than wild-type | In vivo | (19) |

| | | | | |
|---|---|---|---|---|
| **HIV-1 Protease** | | | | |
| N98D | - | N98I was observed when performing transposon-directed base-exchange mutagenesis using a random mutant library, function unknown | In vitro | (20) |
| K70T | - | Minority mutation (2%) associated with resistance to protease inhibitors | Clinical Isolate | (21) |
| **HIV-1 RT p66** | | | | |
| F61S | Finger | F61 plays an important role in strand displacement synthesis, with F61Y and F61L increasing efficiency and reduced processivity, while F61W reducing activity | In vitro | (22) |
| P95L | Palm | P95 is a highly conserved location that makes up important dimerization interface that contributes to formation of bottom of NNRTI pocket | Clinical isolate | (23) |
| | | P95 is a proposed target amino acid in design of novel NNRTIs or dimerization (when together with N137 and P140) | In vitro | (24) |
| A98V | Palm | A98 is a NNRTI-associated mutation (5.2%) | Clinical isolate | (25) |
| | | A98G confers low-level (~2-fold) resistance to NVP with uncertain virological effects which rarely occurs in drug-naive patients.<br>A98S is a common polymorphism not associated with NNRTI resistance. | Clinical isolate | (26) |
| | | A98G is associated with etravirine resistance, polymorphic in non-B subtypes | Clinical isolate | (27, 28) |
| | | A98S is present at low variability in drug-naïve patient (6.8%) | Clinical isolate | (23) |
| | | A98G is selected by nevirapine (NVP) | Clinical Isolate; In vitro | (29) |
| K103R | Palm | No changes in NNRTI susceptibility alone. Has synergistic effect on NNRTI when combined with V179D | Clinical Isolate; In vitro | (30) |
| V108M | Palm | V108 is a NNRTI-associated mutation (15.2%) | Clinical Isolate | (31) |
| | | V108I shown to indirectly confer resistance via alterations of drug stacking interactions of the drug through Y181 | Crystallography | (32) |
| N136D | Finger | N136 is essential to preserve catalytic activity, resulting in increased amounts of free p51 and p66 monomers. Mutant N136D decreased inhibitory activity of NNRTI by 1.4- to 6-fold. | In vitro | (33) |
| E204D | Palm | E204D and E204K are polymorphisms observed in drug naïve patients | Clinical Isolate | (34) |
| L214P | Palm | L214K found in 6 patients who failed rilpivirine-containing ART | Clinical Isolate | (35) |
| L228P | Thumb | L228I confers low-level resistance to etravirine, and in combination with Y188C, displays high level of cross resistance to NVP and EFV. | In vitro | (36) |
| | | L228H/R strongly associated with NRTI therapy, L228N is an undifferentiated RTI-selected mutation | Clinical Isolate | (37) |

| | | | | |
|---|---|---|---|---|
| | | L228 substitutions strongly associated with TAMs in treated patients | Clinical Isolate | (38) |
| | | L228Q associated with NVP resistance in NRTI-exposed, NNRTI-naive subjects | Clinical Isolate; In vitro | (39) |
| | | L228H/M/R is a polymorphism associated with reduced virological response to didanosine (ddI) | Clinical Isolate | (40) |
| | | L228 associated with patients receiving multiple nucleoside analog inhibitors | Clinical Isolate | (41) |
| | | L228H/R involved in regulation of resistance to NNRTIs | Clinical Isolate | (42) |
| W239C | Thumb | W239 interacts through P-P interactions with Y318, involved in resistance to NVP and DLV | Clinical Isolate; In vitro | (43) |
| I326V | Connecti on | Decreased proportion in NRTI-treated patients when compared to treatment-naive | Clinical Isolate | (44) |
| K331R | Connecti on | K331A impairs RT dimerization | In vitro | (45) |
| G333R | Connecti on | G333D/E is critical in facilitating dual resistance of AZT and 3TC resistance mutation | In vitro | (46) |
| N363A | Connecti on | Mutation in p51 subunit reduces ability to associate with p66 unit | In vitro | (47) |
| K366E | Connecti on | K366R is selected in NRTI-treated subjects | Clinical Isolate | (44) |
| T470A | RNase H | Frequency of T470N decreased in treatment-experienced subtype B isolates compared to drug-naïve isolates | Clinical Isolate | (44) |
| | | T470A/N/G/R found in both naïve and pre-treated patients, T470P/S/E/K mutated more frequently in pre-treated patients | Clinical Isolate | (48) |
| V536I | RNase H | Polymorphism | Clinical Isolate | (49) |
| A554D | RNase H | A554T/L/K was found to mutate more frequently in pre-treated than in naive patients, suggesting role in NRTI resistance | Clinical Isolate | (48) |

**Table S6 Change in mutational free energies ($\Delta\Delta G_{Mt}$, kcal/mol) of experimentally generated multiple mutation variants.**

| Gene | Amino acid multiple mutation | $\Delta\Delta G_{Mt}$ (kcal / mol) | | Average $\Delta\Delta G_{Mt}$ (kcal / mol) | |
|---|---|---|---|---|---|
| | | Rosetta Cartesian_ddg | FoldX Build PDB | Rosetta Cartesian_ddg | FoldX BuildPDB |
| Gag | K202R, T204A, I223V, S462R, R214G | 2.29 | 5.65 | 0.37 | 1.75 |
| | E42K,P222L | 1.92 | 0.51 | | |
| | A341S,G396S | 1.61 | 3.31 | | |
| | T470A, T471A | -4.36 | -2.46 | | |
| p66 Wildtype | I326V,K331R,E341G,R353G,N360A,K363E,I375M | 0.80 | 5.09 | 2.53 | 3.25 |
| | K22E,K103R,N136D | -0.56 | -0.37 | | |
| | I380M,N418G,K528E | 6.17 | 6.04 | | |
| | F61S,V518A | 3.69 | 2.24 | | |
| p66 Codon Mutated | R269,T281A,K308E,K351E | 2.63 | 3.01 | 6.48 | 7.00 |
| | T482A,N504D,I507V,K543E,I541V,K543E | 5.81 | 6.07 | | |
| | T482A,N504D,I507V,K525E,I541V,K543E,K332E,Y339C,I367V,K375E,K380R,Y390C,T394A,Y426C,T462A,I467V,N479D,I511V | 12.35 | 16.40 | | |
| | K15R,T24A,K34E,I48V,K49R,K51E,N122D | 6.79 | 1.86 | | |
| | K525E,I527V,N530D,K535R,R542G | 3.09 | 4.34 | | |
| | Y426C,K436R,R448G,K450R,K461G,I467V,T482A,S484G,I491V,N504D,K512G,K515E,K525G,N530D,S538G,K543E | 17.19 | 18.68 | | |
| | T336A,K338R,M342V,K416R,I491V | 0.65 | 2.87 | | |
| | K409E,R433G | 0.37 | 0.24 | | |
| | K416R,Y442C,T444A,Y468C | 9.41 | 9.48 | | |

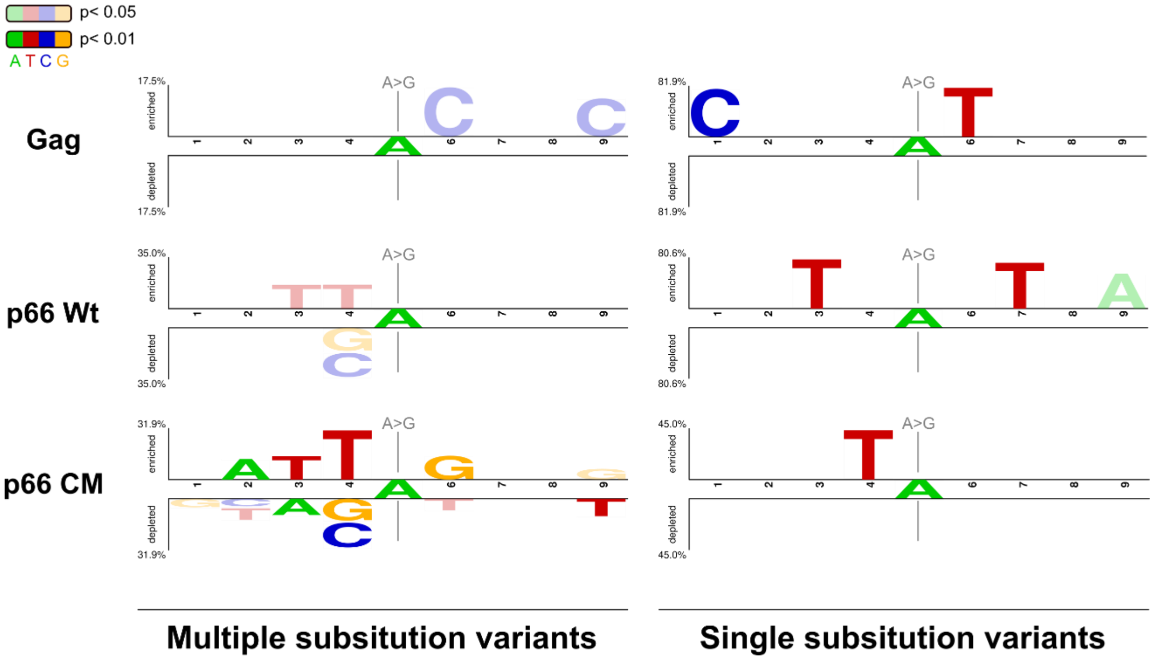


**Fig S1 Two Sample logos of HIV-1 Gag, protease, RT p66 (p66 Wt) and p66 Codon Mutated RT (p66 CM) illustrating the underlying sequence contexts of all adenosine mutations.** Bases were coloured opaque when $p < 0.01$, and translucent when $p < 0.05$.
